# CELL-E 2: Translating Proteins to Pictures and Back with a Bidirectional Text-to-Image Transformer

**DOI:** 10.1101/2023.10.05.561066

**Authors:** Emaad Khwaja, Yun S. Song, Aaron Agarunov, Bo Huang

## Abstract

We present CELL-E 2, a novel bidirectional transformer that can generate images depicting protein subcellular localization from the amino acid sequences (and *vice versa*). Protein localization is a challenging problem that requires integrating sequence and image information, which most existing methods ignore. CELL-E 2 extends the work of CELL-E, not only capturing the spatial complexity of protein localization and produce probability estimates of localization atop a nucleus image, but also being able to generate sequences from images, enabling *de novo* protein design. We train and finetune CELL-E 2 on two large-scale datasets of human proteins. We also demonstrate how to use CELL-E 2 to create hundreds of novel nuclear localization signals (NLS). Results and interactive demos are featured at https://bohuanglab.github.io/CELL-E_2/.

## 1 Introduction

Subcelllular protein localization is a vital aspect of molecular biology as it helps in understanding the functioning of cells and organisms [1]. The correct localization of a protein is critical for its proper functioning, and mislocalization can lead to various diseases [2]. Protein localization prediction models have typically relied on protein sequence data [3, 4] or fluorescent microscopy images [5, 6] as input to predict which subcelleular organelles a protein would localize to, designated as discrete class labels [7, 8]. The CELL-E model was markedly different in that it utilized an autoregressive text-to-image framework to predict subcellular localization as an images [9], thereby overcoming bias from discrete class labels derived from manual annotation [10]. Furthermore, CELL-E was capable of producing a 2D probability density function as an image based on localization data seen throughout the dataset, yielding more a far more interpretable output for the end user.

Although novel, CELL-E was inherently restricted by the following limitations:

### Autoregressive Generation

Alongside other autoregressive models [11–14], CELL-E was limited by slow generation times and unidirectionality. When provided with input, CELL-E required a separate step for each image patch (256 for the output image composed of tokens of size 16 *×* 16). This slow image generation severely limits the ability of CELL-E to perform a high-throughput mutagenesis screening.

### Unidirectional Prediction

The unidirectional nature of CELL-E allowed for predictions to be made in response to an amino acid sequence, however it may be of interest to biologists to make predictions of sequence given a localization pattern. Such capability would be advantageous for those working in a protein engineering domain [15, 16]. One could imagine a researcher wanting to know the optimal localization sequence to append to a protein on either the N or C terminus [17] while maintaining essential function within an active site region, as well as reducing the chance of off-target trafficking.

### Limited Dataset

CELL-E utilized the OpenCell dataset [18], a relatively small dataset. Vision transformers often require large amounts of data to make robust predictions [19], however a small dataset was utilized in the original model. This led to a degree of overfitting and prediction bias based on the limited diversity in localization patterns of the original dataset.

### Present Work

As in CELL-E, our method CELL-E 2 is able to generate accurate protein localization image prediction as illustrated in Fig. 1, but it differs from CELL-E by employing a non-autoregressive (NAR) paradigm which improves the speed of generation. Similar to CELL-E, we retrieve embeddings from a pre-trained protein language model and concatenate these with learned embeddings corresponding to image patch indices coming from a nucleus (a subcellular organelle containing DNA [20]) image and protein threshold image encoders (Fig. 2). We then apply masking to both the amino acid sequence as well as the threshold image in an unsupervised fashion, and reconstructed tokens are predicted in parallel, allowing for generation in fewer steps. This also allows for bidirectional prediction, (sequence to protein threshold image or protein threshold image to sequence). Additionally, to improve the predictive performance we utilize a larger corpus of data, the Human Protein Atlas (HPA) [21] in pre-training to expose the model to a higher degree of localization diversity, and finetune on the OpenCell dataset [18], which better represents natural protein localization because it is acquired from live instead of fixed cells. We explore multiple strategies towards finetuning which serves to generally inform task-specific refinement text-to-image models in Section 5.3. Our code will be made freely available upon publication.

**Figure 1.**
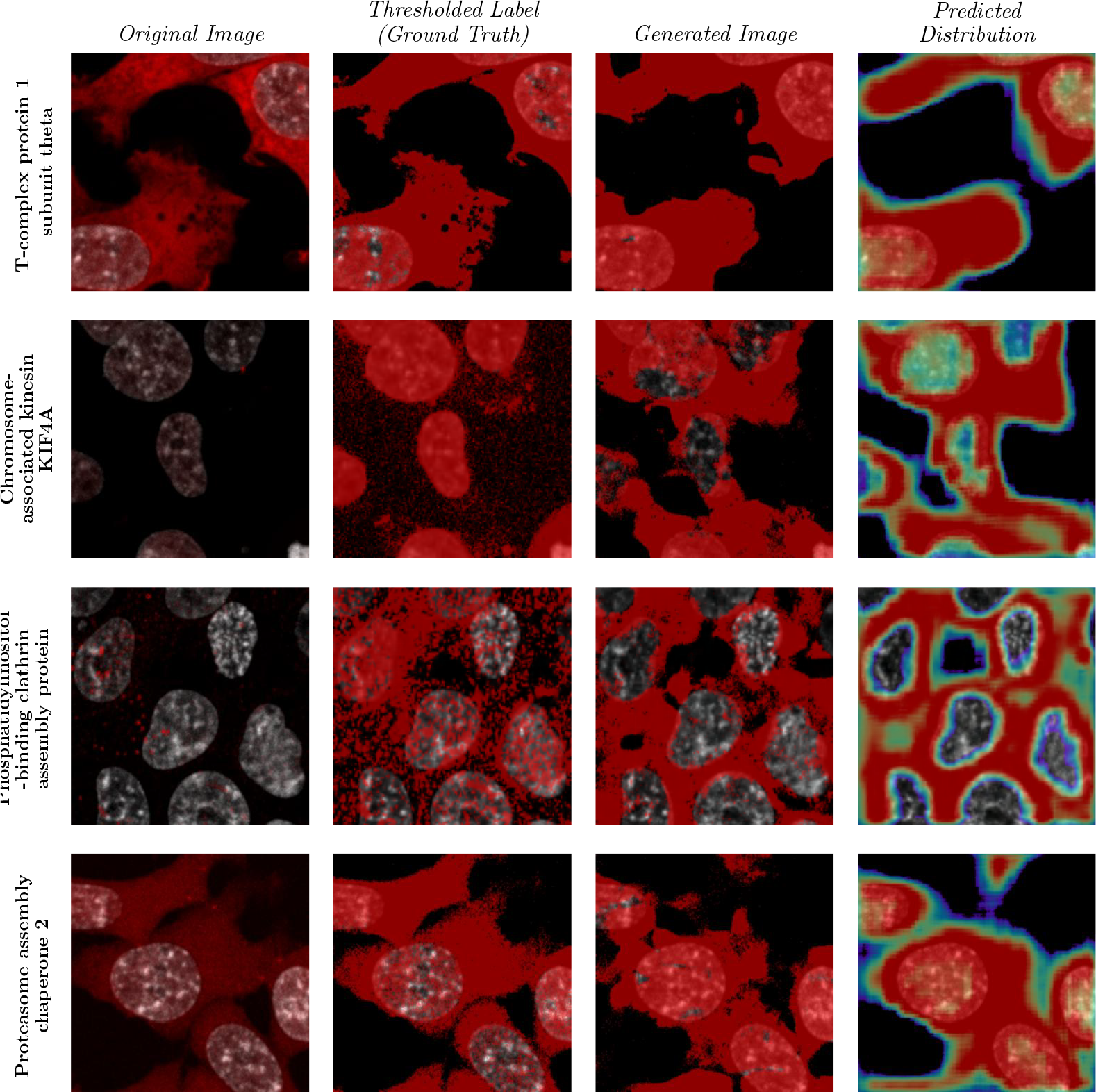
Localization predictions from CELL-E 2 (`HPA Finetuned (Finetuned HPA VQGAN)_480`) on randomly chosen validation set proteins from the OpenCell dataset. All images feature the Hoescht-stained nucleus image as a base. The “Original Image” column shows the fluroscently labelled protein from the dataset. The “Thresholded Label” shows pixels greater than the median value. This serves as the ground truth for the model during training. “Generated Image” is the image specifically predicted by CELL-E 2 and is compared against the thresholded ground truth image. “Predicted Distribution” is the latent space interpolation of the binary threshold image tokens which uses which utilizes the output logits of CELL-E 2. See Fig. S1 for colorbars corresponding to all plots in this work.

**Figure 2.**
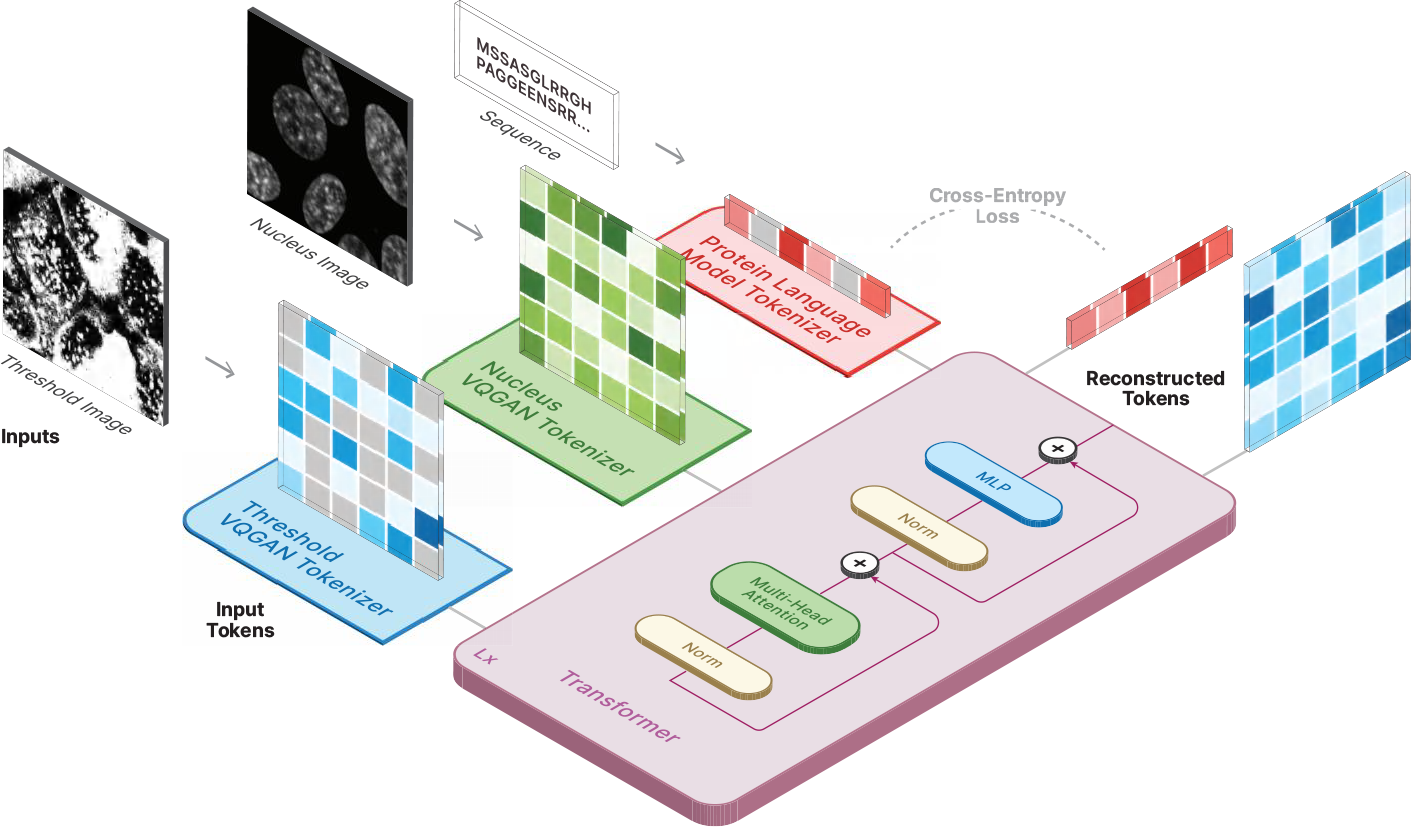
Depiction of training paradigm for CELL-E 2. Gray squares indicate masked tokens. Loss is only calculated on masked tokens in the sequence and protein threshold image.

## 2 Related Work

### 2.1 Protein Language Models

Embeddings from unsupervised protein language models can be used to predict and analyze the properties of proteins, such as their structure, function, and interactions [22]. By exploring the hidden patterns and relationships within these sequences, protein language models can help to advance our understanding of the complex world of proteins and their roles in various biological processes. Masked language modelling has been particularly successful. Ankh [23], ProtT5 [24], ProGen [25], ESM-2 [26], and OmegaFold [27] are examples of recent models which use masked langauge approaches. ESM-2 and Omegafold in particular have been used for structural prediction, indicating hierarchies of information beyond the primary sequence contained in the embeddings [28].

### 2.2 Protein Localization Prediction

Protein localization prediction via machine learning is an emerging field that uses computational algorithms and statistical models to predict the subcellular spatial distribution of proteins [29]. This is an essential task in biology, as the subcellular localization of a protein plays a crucial role in determining its function and interactions with other proteins [30, 31] The prediction of protein localization is performed by analyzing protein sequences, amino acid composition, and other features that can provide clues about their subcellular location. Machine learning algorithms are trained on large datasets of labeled proteins to recognize patterns and make predictions about the subcellular location of a protein. This field has the potential to improve our understanding of cellular processes, drug discovery, and disease diagnosis.

Recently, attention-based methods have demonstrated the ability to predict localization from a sequence [32], enabling the use of long context information when compared to convolutional-based counterparts [33–35]. These methods, however, predict localization as discrete classes rather than as an image. CELL-E, on the contrary, does not utilize existing annotation and provides a heatmap of the expected spatial distribution on a per-pixel basis [9]. This approach enables learning at scale by eliminating the bottleneck of manual annotation while also circumventing label bias.

### 2.3 Text-to-Image Synthesis

Recently, there has been a significant advancement in the field of text-to-image synthesis. Gains have largely been made by autoregressive models [11, 13], which correlate text embeddings with image patch embeddings, as well as diffusion models, [14, 36–39], which condition on sentence embeddings to gradually synthesize images from random noise.

A few works implement non-autoregressive models (NAR), which take advantage of a masked reconstruction procedure, similar to BERT [40], where the model is tasked with predicted randomly masked portions of an input image. These types of models are particulalry advantageous because they enable parallel decoding, allowing images to be synthesized in relatively view steps when compared to autoregressive models. Furthermore, NAR models are not bound to a particular direction of synthesis like autoregressive models, which only perform next-token prediction. CogView2 [41] utilizes a modified transformer architecture where attention on masked tokens is eliminated. MUSE [42] builds on MaskGIT [43] by concatenating a pre-trained text embedding to a token masked representation of a corresponding image. It uses a vanilla transformer architecture [44] and yielded state-of-the-art image synthesis performance in terms of FID and human evaluation.

## 3 Datasets

We pretrained our model on protein images from the Human Protein Atlas (HPA) [45], which covers various cell types and imaging conditions using immunofluorescence microscopy1. We then finetuned on the OpenCell dataset [18], which has a consistent modality using live-cell confocal microscopy of endogenously tagged proteins. To ensure generalization to new data, we followed the homology partitioning method of [35]. We used PSI-CD-HIT [46] to cluster HPA proteins at (≥ 50%) sequence similarity and randomly split the clusters into 80/20 train/validation sets. We applied the same clustering and splitting to the OpenCell proteins, matching the train/validation labels from HPA. For proteins present in OpenCell but not HPA *n* = 176, we assigned the protein based on the other labels in the cluster. Any remaining unassigned proteins *n* = 1 were assigned to the training set. See Appendix A for more details about the datasets and preprocessing.

## 4 Methods

CELL-E 2 (Fig. 2) is a masked *encoder-only* transformer model, which upgrades the capabilities of CELL-E, an autoregressive *decoder-only* model [47]. Due to the NAR nature of the model, CELL-E 2 is capable of both image generation (sequence to image), as well as sequence prediction (image to sequence).

### 4.1 Amino Acid Sequence Embeddings

Proteins are biological molecules which are comprised of individual amino acids. CELL-E 2 utilizes embeddings from ESM-2 [26], where amino acid molecules are denoted with individual letter codes (e.g. A for alanine) [48]. We opt to use frozen embeddings for the prediction task, which has been demonstrated to yield superior reconstruction performance in text-to-image models [9, 37, 42]. The embeddings obtained from a protein language model contain valuable information about amino acid residues, biochemical interactions, structural features, positional arrangements, as well as other characteristics like size and complexity [22]. We train models of varying size based on the released ESM-2 checkpoints (See Section 5). The output of the final embedding layer per respective model is used as the amino acid sequence embedding.

### 4.2 Image Tokenization

Just as in Khwaja et al. [9], we utilize a nucleus image, which serves as a spatial reference with which a binarized protein threshold image is associated. We chose this in order to parallel the type of images which are typicall acquired in a wet lab scenario.

We utilize VQGAN autoencoders [49] trained on both the HPA and OpenCell datasets, respectively. VQGAN surpasses other quantized autoencoders by incorporating a learned discriminator derived from GAN architectures [50]. Specifically, the Nucleus Image Encoder employs VQGAN to represent 256 *×* 256 nucleus reference images as 16 *×* 16 image patches, with a codebook size of (*n* = 512) image patches. To enable transfer learning, we explore finetuning strategies these VQGANs in Section 4.5.

The protein threshold image encoder acquires a compressed representation of a discrete probability density function (PDF) that maps per-pixel protein positions, presented as an image. We binarize the image based on the median pixel value of the image (see Appendix A.4). We utilize a VQGAN architecture identical to the Nucleus VQGAN to estimate the entire set of binarized image patches to denote local protein distributions. These VQGANs are trained until convergence, and the discrete codebook indices are used for the CELL-E 2 transformer. Hyperparameters (Table S1, Table S2, Table S3) and training details can be found in Appendix B.2.

### 4.3 Input Masking Strategy

We adopt a cosine-scheduling technique for masking image tokens, which has been used by other works. The probability of an image patch being masked is determined by a cosine function, favoring high masking rates with an expected masking rate of 64% [42, 43]. This technique provides various levels of masking during the training process, exposing the model to spatial context for masked language tokens.

Of similar interest as image prediction, sequence in-filling with respect to a localization pattern is of interest to biologists. Typically, protein localization sequences are found through sequence similarity searches with proteins that have known localizations in particular organelles [51–53] or via experimentation [54, 55]. CELL-E 2’s bidirectionality enables the model to make predictions for image sequences and sequence predictions for images, making it a novel approach to protein engineering. To achieve this, we also mask the language tokens along with the protein threshold image tokens. We experimented with using the same cosine function for image masking but found it led to numerical instability and vanishing gradients. Therefore, we linearly scaled the cosine function to ensure the maximum masking rate matched 15% masking rate used to train ESM-2.

### 4.4 Base Transformer

The base transformer is an *encoder-only* model in which the embedding dimension is set to the embedding size of the pretrained language model used. We utilized two types of masking tokens. For masking the amino acid sequence, we leveraged the mask token which already exists within the ESM-2 dictionary, designated as `<MASK_SEQ>`. The VQGAN does not contain a masking token within its codebook, so to represent it, we add an additional entry in the image token embedding space of CELL-E 2 (with *n* + 1: (512 + 1 = 513), where *n* is the number of tokens in the VQGAN codebook), and designate the final token as `<MASK_IM>`. We additionally create an embedding space of length 1 for the `<SEP>` token which is appended to the end of the amino acid sequence. Training details can be found in Appendix B.2.

We sample from this transformer by strategically masking positions in the image or sequence (see Appendix B.1). The logit values for the image prediction are used as weights for the threshold image patches to produce a predicted distribution (Fig. 1, Fig. S5).

### 4.5 Finetuning

We sought to leverage useful information from both HPA and OpenCell. HPA contains many proteins (17,268) but is subject to inaccuracies fundamentally because of the immunohistochemistry used for staining, which requires several rounds of fixation and washing [21]. This means the proteins are not observed in a live cell; are subject to signal loss, artifacts, and/or relocalization events; and therefore do not necessarily represent the true nature of protein expression and distribution within a cell [56]. The OpenCell dataset, while comparatively smaller, overcomes these issues by using a split-fluorescent protein fusion system allows for tagging endogenous genomic proteins, maintaining local genomic context, and the preservation of native expression regulation for live cell imaging [18, 57]. We therefore initially trained on HPA, and then finetuned on OpenCell.

Finetuning in the text-to-image domain is still an open question. The use of multiple models makes it difficult to pin down the correct strategy. Contemporary efforts utilize pre-trained checkpoints to fine-tune on domain specific data [58–60]. Chambon et al. [61] reported improved synthesized image fidelity when fine-tuning the U-net of a text-to-image diffusion model, but similar fine-tuning strategies have not been explored for patch-based methods. We report our findings in Section 5.3.

## 5 Results

Similar to CELL-E, we cast the embedding spaces for the image tokens at the same size as the ones used by the pre-trained language model. The size of the embedding vectors (“Hidden Size”) for each model was chosen based on the publicly available ESM-2 checkpoints. For instance, a CELL-E 2 model with hidden size = 480 uses `esm2_t12_35M_UR50D`, which corresponds to a 35M parameter model with 12 attention layers. Khwaja et al. [9] demonstrated a positive relationship between the number of attention layers (designated “Depth”) in the base transformer and the image prediction performance. The maximum depth was set based on our available GPU memory capacity. We refer to models using the name format “`Training Set_Hidden Size`”.

### 5.1 Protein Localization Image Prediction Accuracy

To predict the protein localization image, we provide CELL-E 2 with the protein sequence and nucleus image, and fill the protein image token positions with `<MASK_IM>` tokens (Fig. S3).

We evaluated the models on several image metrics (see Appendix C.1) that measure the quality and diversity of the generated protein images (Table 1). Additionally, we assessed the model’s generalization capabilities by testing them on the other dataset (HPA-trained model on OpenCell and *vice versa*) (Table S4). We report the results for each model on its respective dataset. We observed a significant positive effect of depth on performance across all metrics and datasets. The models with hidden sizes of 480 and 640 achieved the highest scores, with no significant difference between them. However, on the HPA dataset, `HPA_640` surpassed the `HPA_480` model in more categories. On the OpenCell dataset, `OpenCell_480` performed better than the `OpenCell_640`.

**Table 1:** Validation Set Image Prediction Accuracy. MAPE: mean absolute percentage error, MAE: mean absolute error, SSIM: structural similarity index measure, FID: Fréchet inception distance, IS: inception score.

| Dataset | Hidden Size | Depth | Nucleus Proportion MAPE | Image MAE | PDF MAE | SSIM | FID | IS |
| --- | --- | --- | --- | --- | --- | --- | --- | --- |
| HPA | 480 | 68 | **0.0257 ± 0.0250** | 0.3340 ± 0.0788 | 0.2846 ± 0.0985 | 0.2633 ± 0.1781 | 12.0332 | 2.2900 ± 0.0410 |
| | 640 | 55 | 0.0294 ± 0.0278 | **0.3283 ± 0.0805** | **0.2842 ± 0.0991** | **0.2826 ± 0.1827** | 21.7942 | 2.2618 ± 0.0364 |
| | 1280 | 25 | 0.0370 ± 0.0360 | 0.3622 ± 0.0799 | 0.2967 ± 0.0985 | 0.2645 ± 0.1857 | **1.5161** | **2.5440 ± 0.0490** |
| | 2560 | 5 | 0.0818 ± 0.0794 | 0.3516 ± 0.0792 | 0.3104 ± 0.0904 | 0.2558 ± 0.1619 | 23.7977 | 2.1578 ± 0.0290 |
| OpenCell | 480 | 68 | 0.0161 ± 0.0148 | **0.4953 ± 0.0064** | **0.3620 ± 0.1168** | **0.1220 ± 0.1188** | **1.5844** | **2.6069 ± 0.1175** |
| | 640 | 55 | **0.0159 ± 0.0136** | 0.4995 ± 0.0006 | 0.3785 ± 0.1008 | 0.1011 ± 0.1012 | 2.6966 | 2.0974 ± 0.0981 |
| | 1280 | 25 | 0.0272 ± 0.0223 | 0.4996 ± 0.0010 | 0.4359 ± 0.0700 | 0.0694 ± 0.0472 | 8.9102 | 1.3712 ± 0.0432 |
| | 2560 | 5 | 0.0584 ± 0.0511 | 0.4996 ± 0.0005 | 0.4145 ± 0.0889 | 0.0890 ± 0.0667 | 9.5116 | 1.4176 ± 0.0329 |

We also visually inspected some of the generated protein images (Fig. S6, Fig. S7). The output images from the OpenCell models appeared realistic and consistent with the ground truth labels, but they had low entropy in the predicted distribution. This suggests that the models learned to assign high probability to correct tokens, but failed to capture the uncertainty and variability of other valid selections. This could be attributed to the rapid overfitting of the OpenCell models, which limited their generalization ability.

We found that models performed better on their own datasets than on the other dataset. However, the HPA-trained model had higher image prediction performance on the OpenCell dataset than the OpenCell-trained model, with lower PDF MAE values for all categories. The HPA model also had lower FID on the OpenCell validation set, indicating the benefits of having more data despite different imaging conditions. `OpenCell_480` achieved the best scores for 4 out of 8 metrics (MAPE, MAE, SSIM and IS). This performance is likely due to the large number (25 M) of model parameters.

### 5.2 Masked Sequence In-Filling

To test each model’s sequence learning, we used a masked in-filling task similar to the training task. Similar to Section 5.1, we provide CELL-E 2 with a randomly masked (15%) sequence, an unmasked nucleus image, and an unmasked protein threshold image. To select the sequence prediction, we perform a weighted random sampling operation from the 3 amino acids with the highest predicted probabilities. We measured the accuracy as the percentage of correct predictions (noted as “Sequence MAE”, see Appendix C.2). We then embedded each reconstructed sequence with `esm2_t36_3B_UR50D`, the largest model we could fit in memory, with 3B parameters, 36 layers and an embedding dimension of 2560. We computed the mean cosine similarity between the embeddings of the original and reconstructed sequences at masked positions. We show validation results in (Table 2) and all results in (Table S6).

**Table 2:** Validation Set Masked Sequence In-Filling.

| Dataset | Hidden Size | Depth | Sequence MAE | Cosine Similarity |
| --- | --- | --- | --- | --- |
| HPA | 480 | 68 | $0.8628 \pm 0.0951$ | $0.9504 \pm 0.0237$ |
| | 640 | 55 | $0.7917 \pm 0.1245$ | $0.9577 \pm 0.0216$ |
| | 1280 | 25 | $0.6512 \pm 0.1794$ | $0.9708 \pm 0.0163$ |
|  | 2560 | 5 | **$0.5759 \pm 0.2322$** | **$0.9722 \pm 0.0210$** |
| OpenCell | 480 | 68 | $0.7507 \pm 0.1709$ | $0.9533 \pm 0.0285$ |
| | 640 | 55 | $0.6641 \pm 0.1764$ | $0.9610 \pm 0.0272$ |
| | 1280 | 25 | $0.5698 \pm 0.2016$ | $0.9709 \pm 0.0220$ |
|  | 2560 | 5 | **$0.4950 \pm 0.2456$** | **$0.9711 \pm 0.0271$** |

Most models had low performance on this task in terms of reconstruction. This is understandable because the models learned to generate amino acids that were common or frequent in the dataset, but not necessarily correct for the specific sequence. On the other hand, we observed values close to 1 for the cosine similarity, indicating that the predicted amino acids had similar embedding values to the original ones at the masked positions. This could be because the models learned to capture some semantic or structural features of the amino acids, such as polarity or charge, that were reflected in the embedding space and contributed to the biological function of the sequence. Models that used the embedding model with 2560 dimensions had the best performance. For example, `OpenCEll_2560` had the best performance on both metrics, with a MAE of 0.4950 and cosine similarity of 0.9711. When compared to randomly selected amino acids for each position (Table S7), we note significantly higher Sequence MAE and Cosine Similarity.

We also note that the reconstruction ability does not improve the performance of the original language models (Table S5). This may be a result of the combined image/sequence loss used during training or a smaller corpus of data compared to datasets used for the training the original language model.

Evaluation results across both datasets can be found in (Table S6).

### 5.3 Finetuning

While the HPA dataset contains information about a wide variety of proteins, the model does not innately perform as well on the OpenCell data. We considered the potential of utilizing an HPA-trained model and finetune on the OpenCell data, thereby introducing a wider protein context than what is found in the OpenCell data alone while adapting to the imaging conditions and cell type found within the new dataset. We experimented with different finetuning strategies for CELL-E 2 on the OpenCell dataset. We used the pre-trained HPA checkpoint as the starting point for all finetuned models, continuing training on the OpenCell train set. We also evaluated the pre-trained HPA and OpenCell checkpoints without any finetuning as baselines. The finetuned models differed in how they updated the image encoders:

- `HPA Finetuned (HPA VQGAN)`: we kept the original VQGAN image encoders from the HPA checkpoint.
- `HPA Finetuned (OpenCell VQGAN)`: we replaced the image encoders with the Open-Cell VQGANs.
- `HPA Finetuned (Finetuned HPA VQGAN)`: we finetuned the HPA image encoders while keeping the rest of the model frozen, then freeze the image encoders and update the transformer weights.

Fig. S8 shows image predictions on an OpenCell validation protein for models with hidden size = 480. Surprisingly, the pretrained HPA model already achieved strong performance on the OpenCell dataset without any finetuning (see Table S8). The best results were obtained by finetuning both the VQGAN image encoders and using them in the HPA base transformer checkpoint (see Table 3). We attribute the 1.81% improvement in MAE, along with the improvements in FID and IS, to the fine-tuning of both the VQGANs, as it improved the consistency of image patch tokens. This provided the checkpoint with more reliable image patches to generate from. However, swapping the HPA VQ-GAN with an OpenCell one led to a similar losses of distribution information seen in Fig. S7. This could be because the model overfits before being able to learn probabilities across tokens. The learning obstacle comes from the possibility that images patches within the finetuned OpenCell VQGAN have sufficient (or even more) pixel consistency with the images, but the patch positional indices are misaligned with those of the HPA VQGAN. These findings are consistent with those found in analogous text-to-image works utilizing diffusion models.

**Table 3:**
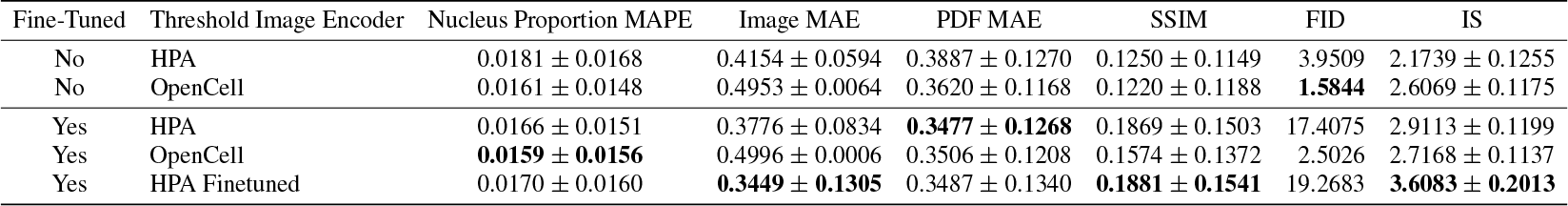
OpenCell Validation Set Image Prediction Accuracy after Finetuning.

| Fine-Tuned | Threshold Image Encoder | Nucleus Proportion MAPE | Image MAE | PDF MAE | SSIM | FID | IS |
| --- | --- | --- | --- | --- | --- | --- | --- |
| No | HPA | 0.0181 ± 0.0168 | 0.4154 ± 0.0594 | 0.3887 ± 0.1270 | 0.1250 ± 0.1149 | 3.9509 | 2.1739 ± 0.1255 |
| No | OpenCell | 0.0161 ± 0.0148 | 0.4953 ± 0.0064 | 0.3620 ± 0.1168 | 0.1220 ± 0.1188 | **1.5844** | 2.6069 ± 0.1175 |
| Yes | HPA | 0.0166 ± 0.0151 | 0.3776 ± 0.0834 | **0.3477 ± 0.1268** | 0.1869 ± 0.1503 | 17.4075 | 2.9113 ± 0.1199 |
| Yes | OpenCell | **0.0159 ± 0.0156** | 0.4996 ± 0.0006 | 0.3506 ± 0.1208 | 0.1574 ± 0.1372 | 2.5026 | 2.7168 ± 0.1137 |
| Yes | HPA Finetuned | 0.0170 ± 0.0160 | **0.3449 ± 0.1305** | 0.3487 ± 0.1340 | **0.1881 ± 0.1541** | 19.2683 | **3.6083 ± 0.2013** |

We did not find that finetuning improved the model’s sequence reconstruction ability (see Table S9).

## 6 Discussion

### 6.1 CELL-E Comparison

In Table S10 and Table S11, we compare the performance of image localization prediction from scratch for CELL-E 2 and CELL-E.

On the OpenCell validation set, CELL-E under-performs CELL-E 2 both before and after finetuning with regards to Nucleus Proportion MAPE. CELL-E 2 achieves worse Image and PDF MAE metrics before finetuning, however after finetuning CELL-E 2 achieves a 2.2% improvement for Image MAE and 1.7% for PDF MAE. On the contrary, CELL-E performs better with respect to image fidelity metrics SSIM and FID.

With respect to speed, we found that the CELL-E 2 with hidden size of 480 was able to generate a prediction 65*×* faster than the CELL-E model. This is a result of CELL-E 2’s capability to generate a prediction in a single step (.2784 seconds). This level of speed enables the advent of large-scale *in silico* mutagenesis studies.

### 6.2 *De novo* NLS Design

CELL-E 2’s bidirectional integration of sequence and image information allows for an entirely novel image-based approach to *de novo* protein design. We applied CELL-E 2 to generate NLSs, which is a short amino acid sequence motif that can relocate a target protein into the cell nucleus when append to the target protein. In this case, our choice of the target protein is the Green Fluorescent Protein (GFP), a common protein engineering target [62–64] that is non-native to the human proteome and absent in the datasets. NLSs are usually identified by experimental mutagenesis studies or *in silico* screens that search for frequent sequences in nuclear proteins [51, 65]. However, these methods may yield candidates that are highly similar to known ones or not specific to the target protein. A more recent approach uses machine learning on sequence identity to augment featurization and statistical priors [17], but it is limited by the distribution of training samples due to the scarcity of experimentally verified NLSs. CELL-E 2 overcomes these limitations because it does not rely on explicit labels, and can therefore leverage significantly more unlabelled image data.

We generated a list of 255 novel NLS sequences for GFP using the procedure described in Appendix D.2. Briefly, we insert mask tokens of set length in a GFP sequence and ask the model with best sequence in-filling performance (`OpenCell_2560`) to fill in the masked amino acids, conditioned on a threshold image generated from the nucleus image (via Cellpose segmentation [66]). To verify the accuracy of the prediction, we pass the predicted sequence through the best performing image model (`HPA Finteuned (Finetuned HPA VQGAN)_480`), and quantify the proportion of signal intensity within the nucleus of the predicted threshold image (Fig. S9). The NLS sequences were then ranked based on sequence and embedding similarity with known NLSs (see Appendix D.2). The list of candidates can be found in Appendix D.3. We found several NLS candidates with high predicted signal in the nucleus, but which were fairly dissimilar from any protein found within NLSdb [65].

Classical NLSs are characterized by having regions of basic, positively charged amino acids arginine (R) and lysine (K) [67, 68], and are categorized as “monopartite” or “bipartite”, either having a single cluster of basic amino acids or two clusters separated by a linker, respectively [69]. We observed a postive correlation between percentage of R and K residues in our predicted NLSs and sequence homology with known NLSs (Table S12). The number of clusters per sequence followed a similar trend, with sequences with relatively low sequence homology (Max ID % ≤ 33) having at most 2 clusters in 88% of predictions (Fig. S10). The remaining predictions, if correct, are therefore non-classical NLSs.

To further verify our predicted sequences, we passed the predicted NLS appended to GFP through Deeploc 2.0 [32], a leading sequence-to-class protein localization model, which predicted 89% of generated sequences were nuclear localizing and 91% contained a potential nuclear localizing signal.

Similar to CELL-E, we observed high attention weights on documented localization sequences correlated with positive protein signal within the threshold image (Fig. S11). For sequences with high predicted nucleus proportion intensities, we observed high activation across the entire sequence (novel NLS and GFP residues), with some NLS weights being an order of magnitude higher than others across the GFP sequences (Fig. 3). On the contrary, predicted sequences with comparatively less predicted intensity within the nucleus had low activation across the sequence, with little to none in the proposed NLS. We observed similar amounts of attention placed on the nucleus image patches, which largely corresponded to the location of the predicted threshold patches

**Figure 3.**
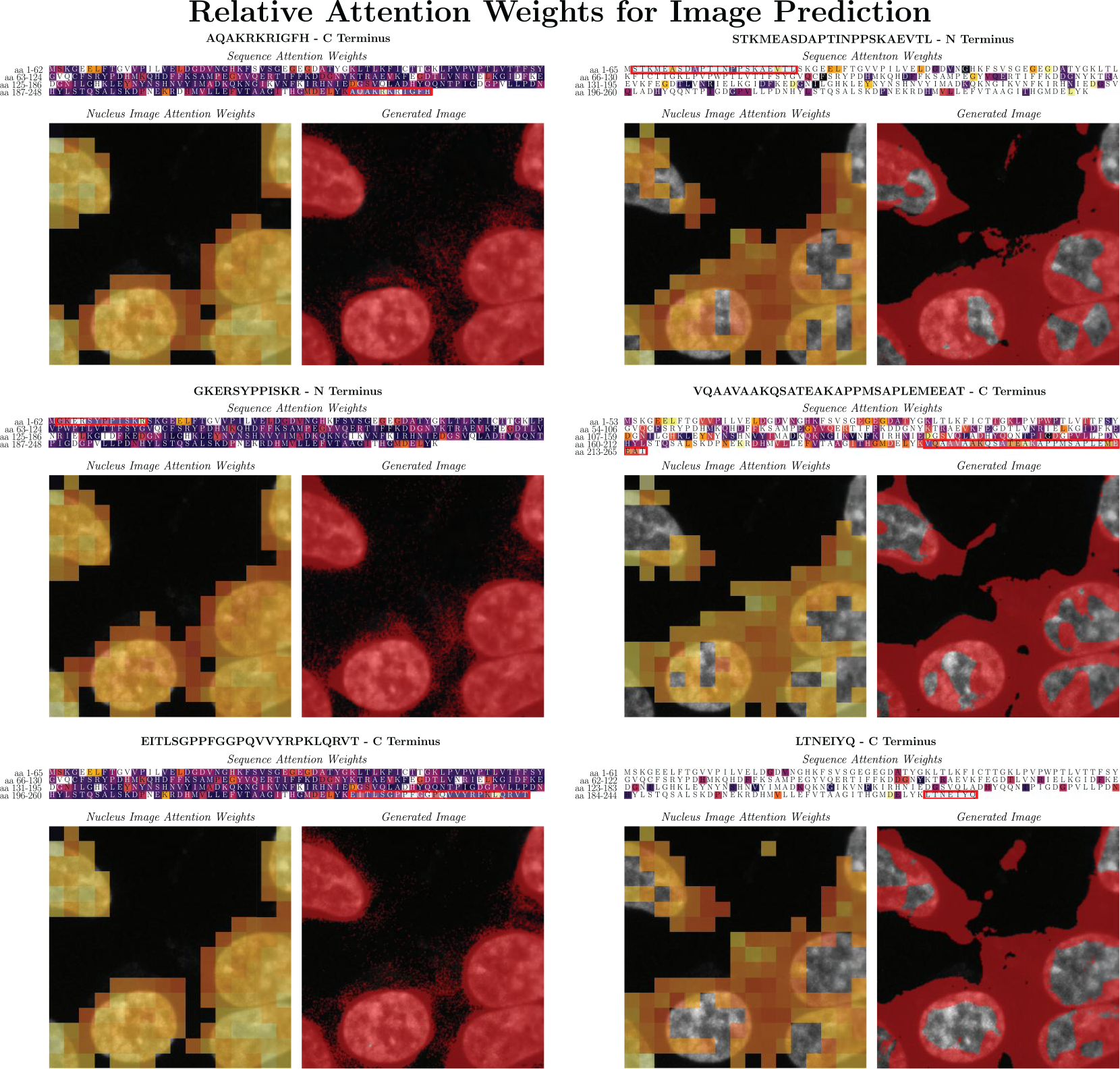
Attention weights associated with positive signal within the predicted image. Tokens with higher attention weight associated with background patches (low signal) are not highlighted. See Appendix D.3 for more information about the visualization process. We show 3 sequences with the highest (left column) and lowest (right column, not included in Table S13) predicted nucleus proportion intensity. The NLS+GFP sequences are shown with the designed NLS boxed in red.

## 7 Conclusion & Future Work

In this paper, we have presented CELL-E 2, a novel bidirectional NAR model for protein design and engineering. CELL-E 2 can generate both image and sequence predictions, handle multimodal inputs and outputs, and run significantly faster than the state-of-the-art. By pre-training on a large HPA dataset and fine-tuning on OpenCell, CELL-E 2 can achieve competitive or superior performance on image and sequence reconstruction tasks. However, one limitation of CELL-E 2 is its output resolution, which is currently (256 *×* 256). This resolution may not capture the fine details of microscopy images, which may limit applications in real-world use where Megapixel images are acquired. Increasing the output resolution of CELL-E 2 is one direction for future work. Furthermore, the sequence prediction struggles with the prediction of large stretches of amino acids as opposed to singular masked positions. Within this work, we encountered a trade-off between sequence prediction quality and prediction speed which may be overcome by reformulating the masking strategy. Similar findings were seen when compared with CELL-E, where we found accuracy measurements to improve with CELL-E 2 at the detriment of image quality metrics. The in-order prediction sequence we utilized in this paper may serve as a bottleneck for protein engineering applications despite the speed advantages gained from using a NAR architecture.

Another direction for future work is to incorporate structural information into the sequence embeddings. CELL-E 2 can generate novel NLS sequences with similar properties to GFP but low homology to existing sequences. However, the current sequence embeddings are based on a language model that may not capture all the structural features of the proteins. These features may affect the image appearance and vice versa.

We believe that CELL-E 2 is a promising model for protein design and engineering. We hope that our work will inspire more research on bidirectional NAR models for this domain and other domains that involve multimodal data.

## Supporting information

List of CELL-E 2 generated NLS sequences

## 8 Acknowledgments

This research is supported in part by NIH grants R01GM131641 (BH) and R35-GM134922 (YSS). B.H. is a Chan Zuckerberg Biohub Investigator. A.A. contributed to visualizations and demos.

## Supplementary Material

### A Datasets

**Figure S1:**
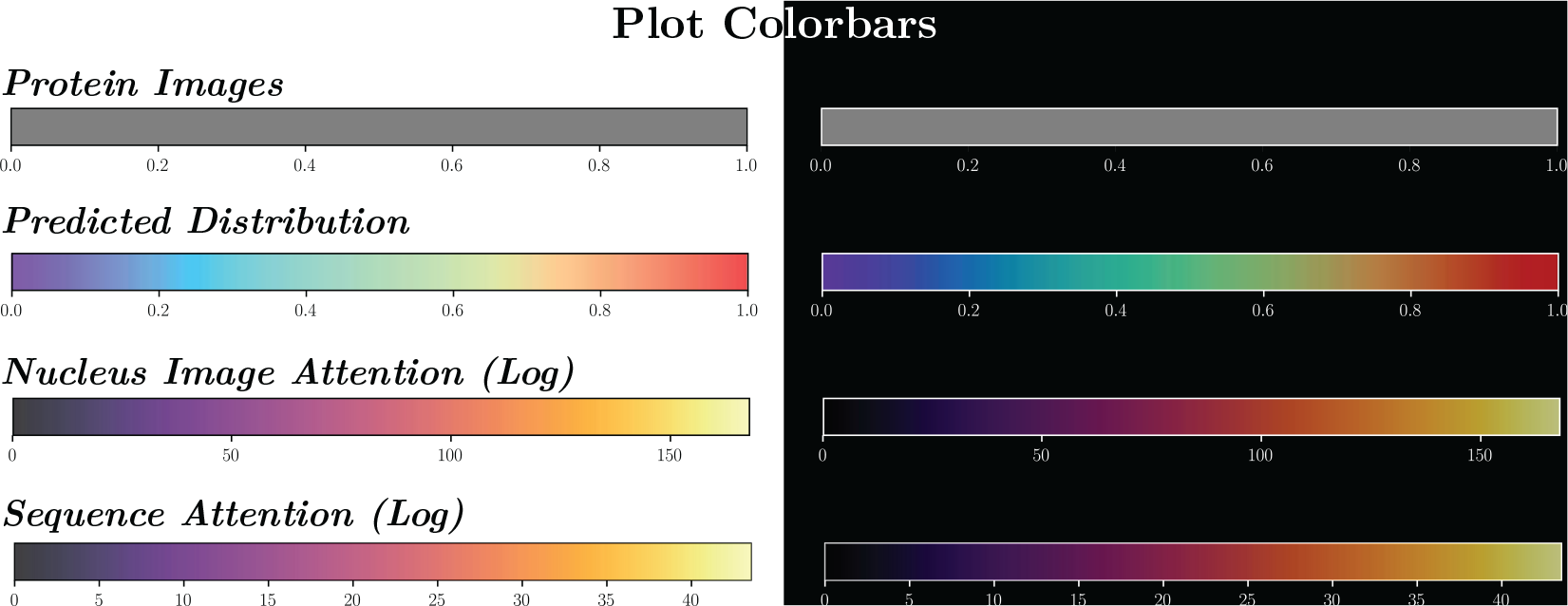
Colorbars used in figures on white (left) and black (right) background.

#### A.1 Human Protein Atlas

We used the Human Protein Atlas v21, available under the Creative Commons Attribution-ShareAlike 3.0 International License. For pre-training, we selected the immunofluorescence stained images from the Human Protein Atlas (HPA), which contains data on more than 17,268 human proteins, with information on their distribution across 44 different normal human tissues and 20 different cancer types. Example images show distribution of proteins within 2-5 cell types with different antibody markers [1]. We extracted corresponding amino acid sequences from UniProt [2].

#### A.2 OpenCell

We selected the OpenCell dataset for fine tuning due to its high-quality images, consistent imaging and cell conditions, and availability of reference images with consistent morphology. The dataset includes a collection of 1,311 CRISPR-edited HEK293T human cell lines, each tagged with a target protein using the split-mNeonGreen2 system. For each cell line, the OpenCell imaging dataset contains 4-5 confocal images of the tagged protein, accompanied by DNA staining to serve as a reference for nuclei morphology. While smaller in comparison to HPA, the cells were imaged while alive, providing a more accurate representation of protein distribution within the cell than immunofluorescence [3]. The OpenCell dataset is available under the BSD 3-Clause License.

#### A.3 Amino Acid Sequence Preprocessing

In natural language contexts, ensuring input sequences are the same length is usually performed by modifying the end of the sequence, either via truncation or end-padding [4]. This allows for predictions with respect to a given input (i.e. a text prompt). From the perspective of protein function, however, both the beginning and end (N and C termini) are points of interest for appending amino acids, especially with respect to protein localization [5, 6]. As such, we augment the sequence data as follows:

1. The amino acid sequence is tokenized using the ESM-2 tokenizer.
2. Start and end tokens are appended to the beginning and end of the sequence.
3. Cropping or padding occur based on the full sequence length, (length of amino acid sequence + <START> token + `<END>` token = 1002).
  - If the full sequence length *>* 1002 tokens, we randomly crop 1002 tokens.
  - If the full sequence length *<* 1002 tokens, we randomly add pad tokens before the `<START>` token and/or after the `<END>` token (See Fig. S2).
4. A `<SEP>` token is appended to the end of the protein sequence.

**Figure S2:**
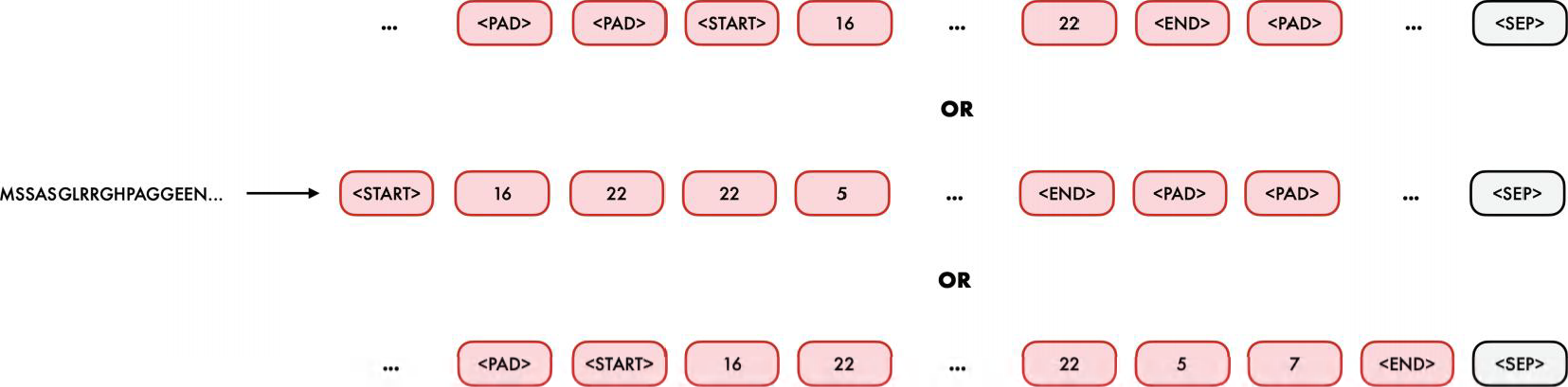
The amino acid sequence is tokenized and randomly padded via the `<PAD>` token. The top row shows start and end padding. The middle row shows end padding. The bottom row shows start-padding. All of these are possible. Note that the fixed length of 1002 means that the `<SEP>` token is always placed in the 1003rd position.

#### A.4 Image Preprocessing

A few preprocessing steps were necessary for the image encoder. Our image processing procedure is as follows:

1. We clip pixels beneath the .001 and above the 99.999 percentiles.
2. We normalize image values based on the calculated means and standard deviation from the datasets: **Human Protein Atlas** Nucleus: *μ* = 0.0655, *σ* = 0.1732 Protein Image: *μ* = 0.0650, *σ* = 0.1208 **OpenCell** Nucleus: *μ* = 0.0272, *σ* = 0.0486 Protein Image: *μ* = 0.0244, *σ* = 0.0671
3. We rescale the images so pixel values are between 0 and 1.
4. The median pixel value of the protein image is calculated to create the thresholded image such that pixels ≥ median = 1 and pixels *<* median = 0. Finally, we rescale images to 600 *×* 600 and randomly crop to 256 *×* 256 pixels.
5. Data augmentation is applied via random horizontal and vertical flips.

## B Methods

### B.1 Sampling

We experimented with the cosine-scheduling approach used in other works [7, 8], but we did not see any improvement in reconstruction performance (Fig. S4). We predicted the entire image in one step for image prediction. For amino acid sequence prediction, we predict amino acids one-by-one from the central protein.

We also calculated the probabilities of each token for all image predictions. We kept the output logits of the transformer. For image logits, we normalized them to 1 and fed them to the VQGAN decoder, which performed a linear interpolation in latent space. We clipped the values between 0 and 1 and displayed them as a heatmap (Fig. S3).

### B.2 Training

We utilized 4*×* NVIDIA RTX 3090 TURBO 24G GPUs for this study. 2 GPUs were utilized for training VQGANs via distributed training. Our computer also contained 2*×* Intel Xeon Silver and 8*×* 32768mb 2933MHz DR *×* 4 Registered ECC DDR4 RAM. Only a single GPU is ever used to train CELL-E 2 models. Models were implemented in Python 3.11 using Pytorch 2.0 [9].

**Figure S3:**
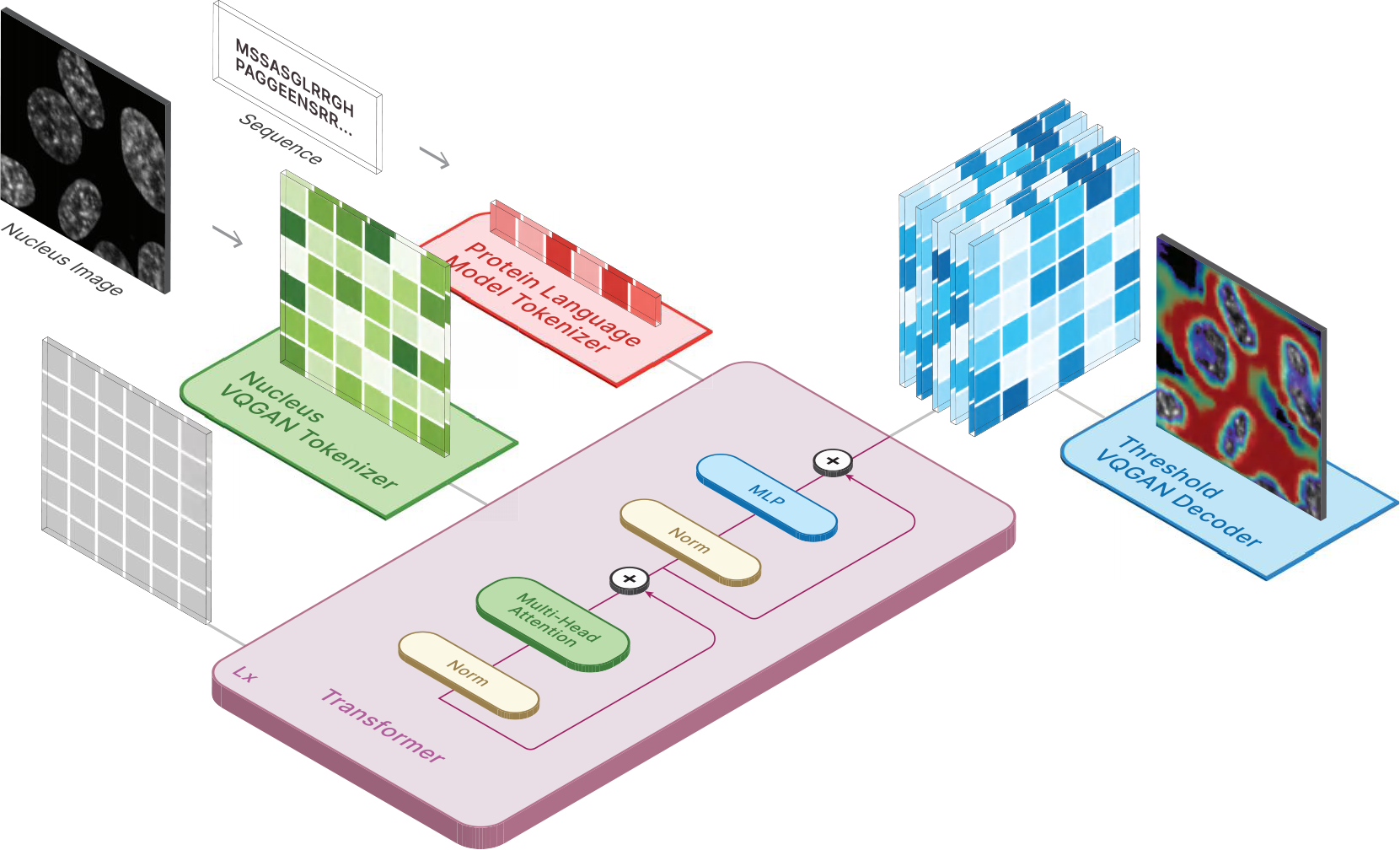
Depiction of the reconstruction scheme used to generate the predicted distribution heatmaps. Similar to training time, we provide tokenized vectors corresponding to the amino acid sequence and the nucleus image. Every position for the tokenized image is set to `<MASK_IM>` (shown as gray squares). The output logits are saved for every position and treated as probabilities associated with each image patch. These values are scaled and sent to the threshold VQGAN decoder to produce the final heatmap. Pixel values in the final image are clipped between 0 and 1.

In order to train the transformer, we underwent the following procedure (Fig. 2):

1. We tokenize the amino acid sequence using the ESM-2 dictionary. We tokenize the nucleus image and protein threshold image using the codebook indices of the respective pre-trained VQGANs.
2. We retrieve embeddings for the amino acid sequence from the pre-trained ESM-2 protein language model (available under the MIT license Copyright (c) Meta Platforms, Inc. and affiliates.). These embeddings are frozen and never updated over the course of training.
3. We randomly mask the amino acid sequence and protein threshold image tokens. The `<SEP>` and nucleus image tokens are never masked.
4. We obtain embeddings for the image tokens from embedding spaces created within the transformer and are learned over training. These size of the embedding are set to the same dimension as the pre-trained language embeddings. We similarly retrieve embeddings from a separate embedding space for the `<SEP>` token.
5. We pass the embeddings through a positional encoder via rotary encoding [10].
6. We concatenate the embeddings along the sequence dimension and pass them through the transformer. We calculate loss via cross-entropy only on the masked tokens.

#### Hyperparameters

We used the following hyperparameters for our transformer model. Based on the findings of Khwaja et al. [11], we increased the transformer depth to achieve better predictive performance. The “Embedding Dimension” was determined by the protein language model we used, so we maximized the number of layers within the transformer, constrained by the VRAM capacity.

**Figure S4:**
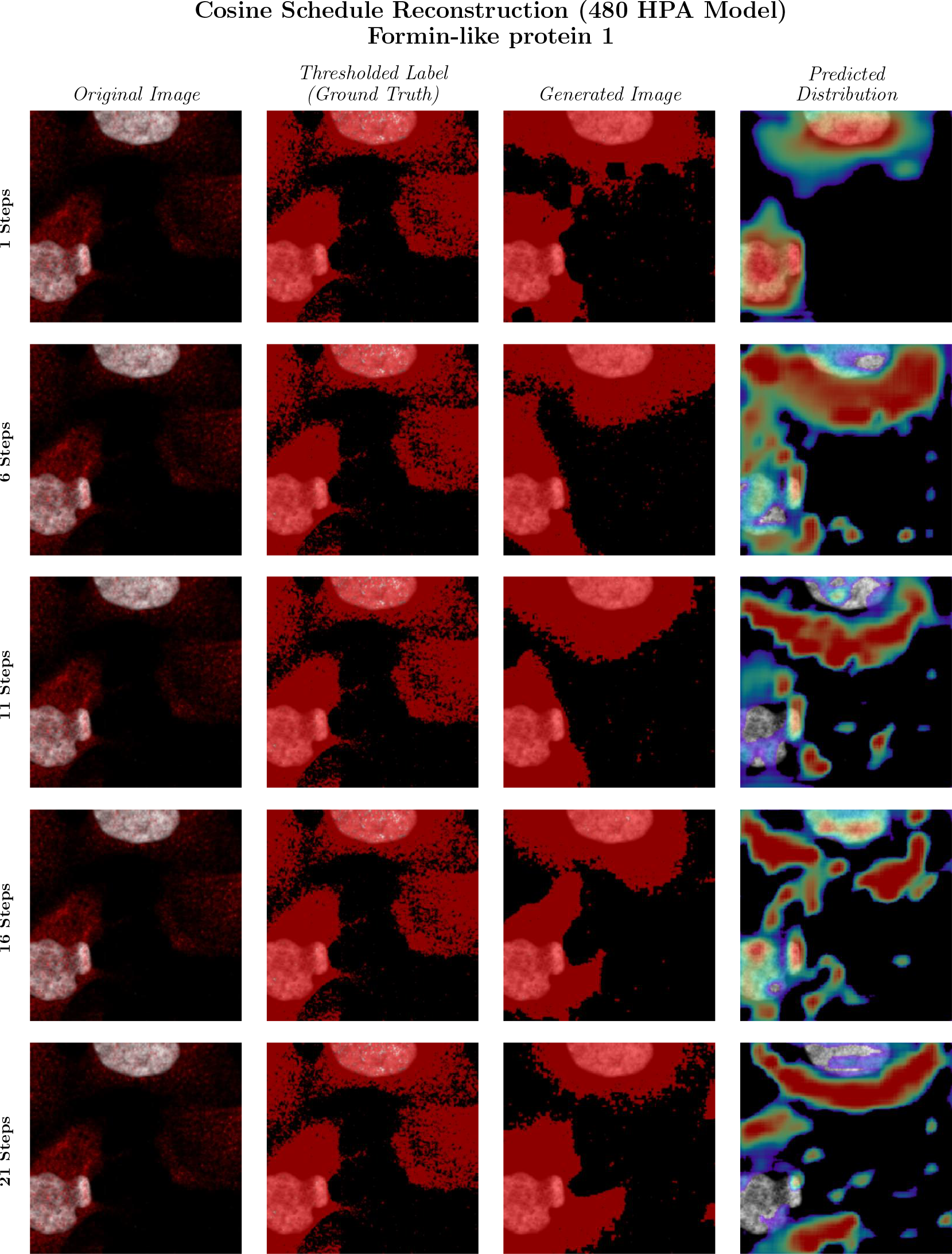
Image prediction based on the number of reconstruction steps. Note the decreased distribution intensity with increasing step count.

**Table S1:** VQGAN Hyperparameters.

| Hyperparameter | Value |
| --- | --- |
| Optimizer | Adam [12] |
| Base Learning Rate | $4.5 \times 10^{-6}$ |
| Betas | $\beta_1 = .5, \beta_2 = .9$ |
| Weight Decay | 0 |
| Embedding Dimension | 256 |
| Number of Embeddings | 512 |
| Resolution | 256 |
| Number of Input Channels | 1 |
| Dropout | 0 |
| Discriminator Start | 50000 |
| Discriminator Weight | .2 |
| Codebook Weight | 1.0 |

**Table S2:** Base Transformer Hyperparameters.

| Hyperparameter | Value |
| --- | --- |
| Optimizer | AdamW [13] |
| Base Learning Rate | $3 \times 10^{-4}$ |
| Betas | $\beta_1 = .9, \beta_2 = .95$ |
| Weight Decay | .01 |
| Number of Text Tokens | 33 |
| Text Sequence Length | 1000 |
| Embedding Dimension/Depth | 480/68 or 640/55<br>or 1280/25 or 2560/5 |
| Number of Heads | 16 |
| Dimension of Head | 64 |
| Attention Dropout | .1 |
| Feedforward Dropout | .1 |
| Image Loss Weight | 1 |
| Condition Loss Weight | 1 |

**Table S3:**
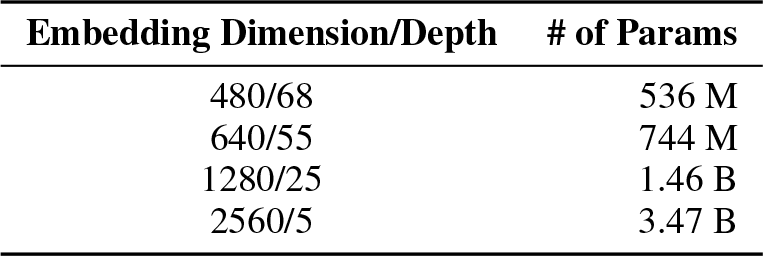
CELL-E 2 Model Parameters per Size.

| Embedding Dimension/Depth | # of Params |
| --- | --- |
| 480/68 | 536 M |
| 640/55 | 744 M |
| 1280/25 | 1.46 B |
| 2560/5 | 3.47 B |

## C Results

### C.1 Image Prediction Accuracy

Table S4 shows the image prediction performance of HPA and OpenCell-trained across both datasets and splits. We evaluate image reconstruction using the following metrics:

#### Nucleus Proportion MAPE

This metric measures how well the predicted protein image matches the ground truth in terms of the fraction of intensity within the nucleus. We use Cellpose [14] to create a mask of the nucleus channel. Then we divide the sum of the predicted 2D PDF pixels inside the mask by the sum of all pixels in the image. We do the same for the ground truth protein image and compare the two fractions. The error is expressed as a percentage of the ground truth fraction.

#### Image MAE

This metric calculates the average absolute difference between each pixel in the predicted protein threshold image and the ground truth protein threshold image. A lower MAE means a better match.

**Figure S5:**
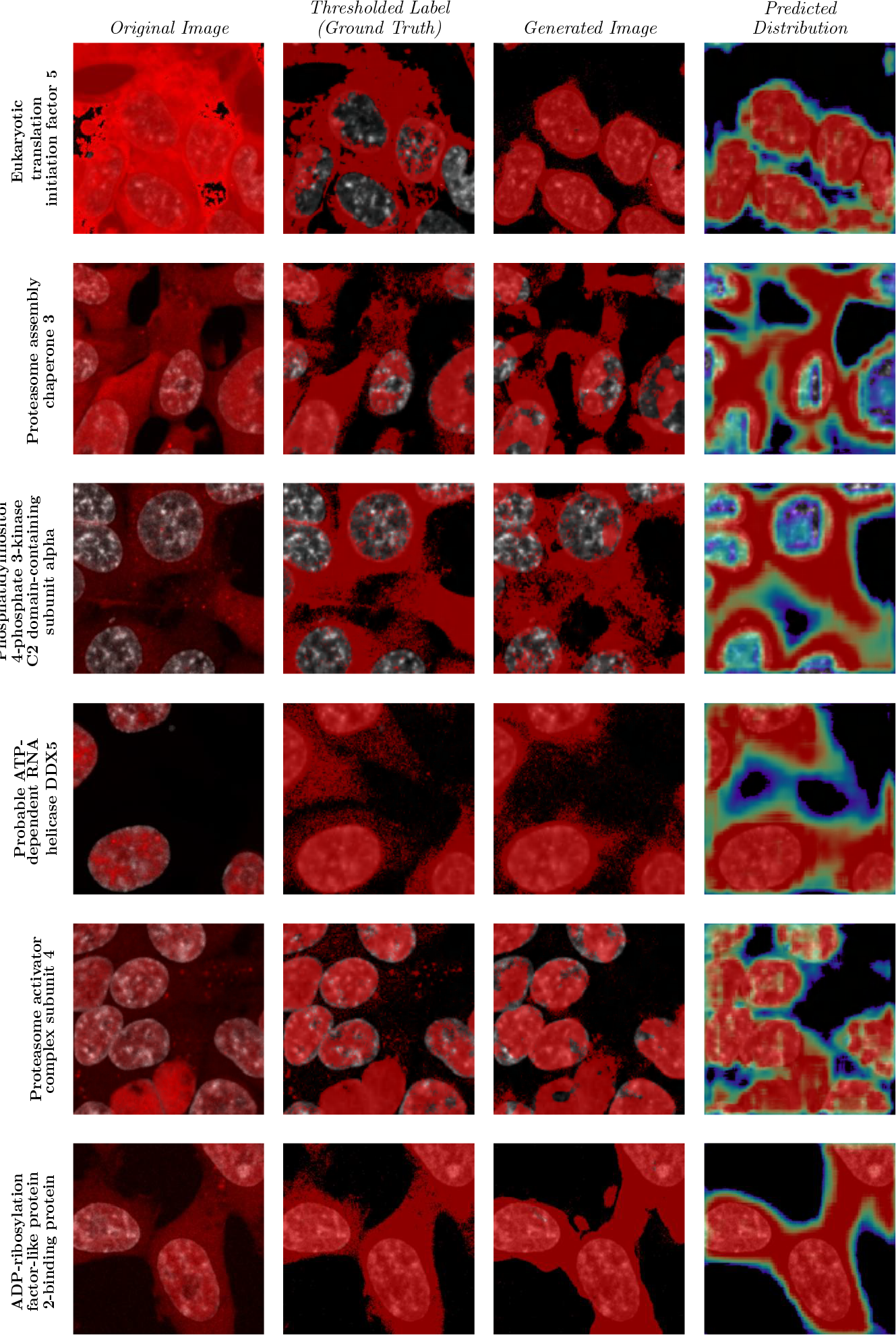
More randomly selected predictions from `HPA Finetuned HPA VQGAN_480`. We only note an incorrect prediction in Eukaryotic translation initiation factor 5.

#### PDF MAE

This metric is similar to Image MAE, except we evaluate the difference using the predicted 2D PDF, rather than the predicted protein threshold image. We expect this number to be less accurate as tokens with less confidence will reduce the pixel value, while all values in the protein threshold image are 0 or 1.

#### SSIM

Structural similarity index measure (SSIM) is a metric that evaluates how similar two images are in terms of local features such as brightness and contrast. It takes into account the spatial relationships between neighboring pixels. SSIM values range from 0, meaning no similarity, to 1, meaning perfect similarity.

**Table S4:** Image Prediction Accuracy Across OpenCell and HPA.

| Training Set Proteins |  |  |  |  |  |  |  |  |  |
| --- | --- | --- | --- | --- | --- | --- | --- | --- | --- |
| Dataset | Train Set | Hidden Size | Depth | Nucleus Proportion MAPE | Image MAE | PDF MAE | SSIM | FID | IS |
| HPA | HPA | 480 | 68 | .0254 ± .0296 | .3344 ± .0797 | .2845 ± .0991 | .2635 ± .1797 | 11.4596 | 2.3151 ± .0224 |
|  |  | 640 | 55 | .0291 ± .0318 | **.3286 ± .0808** | **.2843 ± .0996** | **.2827 ± .1836** | 21.0591 | 2.2879 ± .0153 |
|  |  | 1280 | 25 | .0356 ± .0341 | .3640 ± .0797 | .2942 ± .0973 | .2673 ± .1862 | **1.0080** | 2.5634 ± .0192 |
|  |  | 2560 | 5 | .0788 ± .0773 | .3530 ± .0795 | .3097 ± .0904 | .2569 ± .1636 | 22.8721 | 2.1817 ± .0166 |
|  | OpenCell | 480 | 68 | **.0244 ± .0317** | .4620 ± .0769 | .3530 ± .0803 | .0865 ± .0714 | 4.1290 | **2.7063 ± .0146** |
|  |  | 640 | 55 | .0247 ± .0285 | .4676 ± .0778 | .3572 ± .0781 | .0800 ± .0674 | 37.6196 | 2.4858 ± .0169 |
|  |  | 1280 | 25 | .0368 ± .0321 | .4678 ± .0776 | .3835 ± .0659 | .0712 ± .0518 | 21.3462 | 1.5207 ± .0020 |
|  |  | 2560 | 5 | .0706 ± .0737 | .4678 ± .0777 | .3474 ± .0797 | .1041 ± .0725 | 14.4177 | 1.7531 ± .0109 |
| OpenCell | HPA | 480 | 68 | .0184 ± .0177 | .4138 ± .0573 | .3699 ± .1262 | .1388 ± .1206 | 3.7217 | 2.3090 ± .0548 |
|  |  | 640 | 55 | .0183 ± .0166 | **.4087 ± .0579** | .3835 ± .1191 | .1230 ± .1128 | 3.5440 | 2.0354 ± .0998 |
|  |  | 1280 | 25 | .0219 ± .0202 | .4358 ± .0588 | .3659 ± .1141 | .1225 ± .1198 | 7.1451 | 2.1888 ± .0776 |
|  |  | 2560 | 5 | .0460 ± .0418 | .4164 ± .0693 | .3905 ± .0962 | .0984 ± .0870 | 7.5480 | 2.0104 ± .0519 |
|  | OpenCell | 480 | 68 | **.0134 ± .0131** | .4930 ± .0074 | **.3264 ± .1108** | **.1620 ± .1429** | **.8923** | **3.0345 ± .1000** |
|  |  | 640 | 55 | .0141 ± .0124 | .4994 ± .0006 | .3473 ± .0995 | .1291 ± .1195 | 2.8314 | 2.3160 ± .0702 |
|  |  | 1280 | 25 | .0277 ± .0230 | .4996 ± .0007 | .4276 ± .0707 | .0743 ± .0518 | 9.3420 | 1.3759 ± .0213 |
|  |  | 2560 | 5 | .0567 ± .0479 | .4996 ± .0006 | .4037 ± .0877 | .0927 ± .0681 | 9.8328 | 1.4463 ± .0260 |

| Validation Set Proteins |  |  |  |  |  |  |  |  |  |
| --- | --- | --- | --- | --- | --- | --- | --- | --- | --- |
| Dataset | Train Set | Hidden Size | Depth | Nucleus Proportion MAPE | Image MAE | PDF MAE | SSIM | FID | IS |
| HPA | HPA | 480 | 68 | .0257 ± .0250 | .3340 ± .0788 | .2846 ± .0985 | .2633 ± .1781 | 12.0332 | 2.2900 ± .0410 |
|  |  | 640 | 55 | .0294 ± .0278 | **.3283 ± .0805** | **.2842 ± .0991** | **.2826 ± .1827** | 21.7942 | 2.2618 ± .0364 |
|  |  | 1280 | 25 | .0370 ± .0360 | .3622 ± .0799 | .2967 ± .0985 | .2645 ± .1857 | **1.5161** | 2.5440 ± .0490 |
|  |  | 2560 | 5 | .0818 ± .0794 | .3516 ± .0792 | .3104 ± .0904 | .2558 ± .1619 | 23.7977 | 2.1578 ± .0290 |
|  | OpenCell | 480 | 68 | **.0245 ± .0235** | .4622 ± .0767 | .3533 ± .0803 | .0861 ± .0718 | 41.5344 | **2.6712 ± .0225** |
|  |  | 640 | 55 | .0248 ± .0231 | .4676 ± .0776 | .3575 ± .0783 | .0795 ± .0681 | 38.3386 | 2.4850 ± .0381 |
|  |  | 1280 | 25 | .0371 ± .0343 | .4678 ± .0775 | .3833 ± .0661 | .0713 ± .0525 | 21.6973 | 1.5206 ± .0152 |
|  |  | 2560 | 5 | .0717 ± .0722 | .4678 ± .0776 | .3474 ± .0796 | .1038 ± .0731 | 14.7231 | 1.7524 ± .0160 |
| OpenCell | HPA | 480 | 68 | .0181 ± .0168 | .4154 ± .0594 | .3887 ± .1270 | .1250 ± .1149 | 3.9509 | 2.1739 ± .1255 |
|  |  | 640 | 55 | .0178 ± .0165 | **.4058 ± .0574** | .3651 ± .1197 | **.1359 ± .1183** | 3.0867 | 2.1508 ± .0384 |
|  |  | 1280 | 25 | .0227 ± .0213 | .4323 ± .0581 | .3886 ± .1128 | .1051 ± .1140 | **1.4713** | 2.0247 ± .1003 |
|  |  | 2560 | 5 | .0487 ± .0453 | .4202 ± .0722 | .4049 ± .0870 | .0874 ± .0792 | 9.1799 | 1.9269 ± .0768 |
|  | OpenCell | 480 | 68 | .0161 ± .0148 | .4953 ± .0064 | **.3620 ± .1168** | .1220 ± .1188 | 1.5844 | **2.6069 ± .1175** |
|  |  | 640 | 55 | **.0159 ± .0136** | .4995 ± .0006 | .3785 ± .1008 | .1011 ± .1012 | 2.6966 | 2.0974 ± .0981 |
|  |  | 1280 | 25 | .0272 ± .0223 | .4996 ± .0010 | .4359 ± .0700 | .0694 ± .0472 | 8.9102 | 1.3712 ± .0432 |
|  |  | 2560 | 5 | .0584 ± .0511 | .4996 ± .0005 | .4145 ± .0889 | .0890 ± .0667 | 9.5116 | 1.4176 ± .0329 |

#### IS

Inception score (IS) is a metric that assesses how realistic and diverse the images generated by a model are. It uses a pretrained neural network to classify the images and computes a score based on how well they fit into different categories. A higher IS means more realistic and varied images.

#### FID

Fréchet Inception Distance (FID) is another metric that compares the quality and diversity of generated images to ground truth images. It calculates the distance between two statistical representations of the image distributions, called feature vectors, which are extracted by a pretrained neural network. A lower FID means more similar distributions and better quality images. For this study FID was scored against the training or validation sets when applicable.

### C.2 Masked Sequence In-Filling

Table S6 shows the sequence prediction performance (predicting 15% of masked residues) of the models shown in Table S4. We evaluate only on masked positions using the following criteria:

#### Sequence MAE

This metric calculates the average absolute difference between each amino acid in the predicted sequence and the ground truth sequence for the masked positions. A lower MAE means a better match.

#### Cosine Similarity

We evaluate cosine similarity of the amino acid embeddings. This metric measures the angle between two vectors that represent the predicted sequence and the ground truth sequence. It ranges from -1 to 1, where 1 means the vectors are identical, 0 means they are orthogonal, and -1 means they are opposite. A higher cosine similarity means a more similar sequence. Note that cosine similarity is performed on the entirety of the protein and not just masked positions.

**Figure S6:**
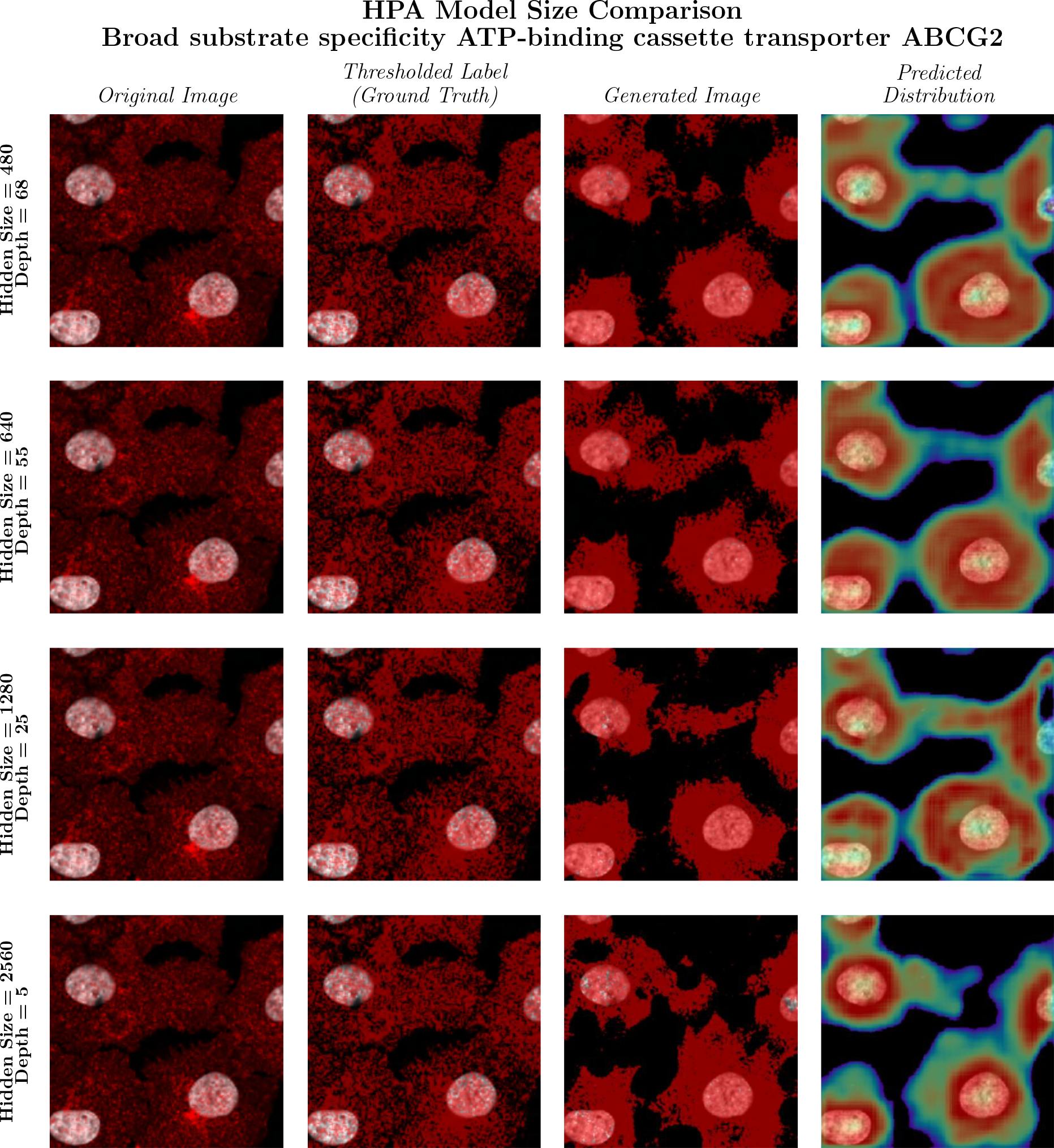
CELL-E 2 models trained on the HPA dataset. Predictions are shown based on the hidden size of the transformer embedding. We see the strongest performance from the 480 and 640 models. Localization is expected within the mitochondria in the selected protein. Not the heightened intensity within the nuclear region in the 1280 and 2560 models predictions.

**Figure S7:**
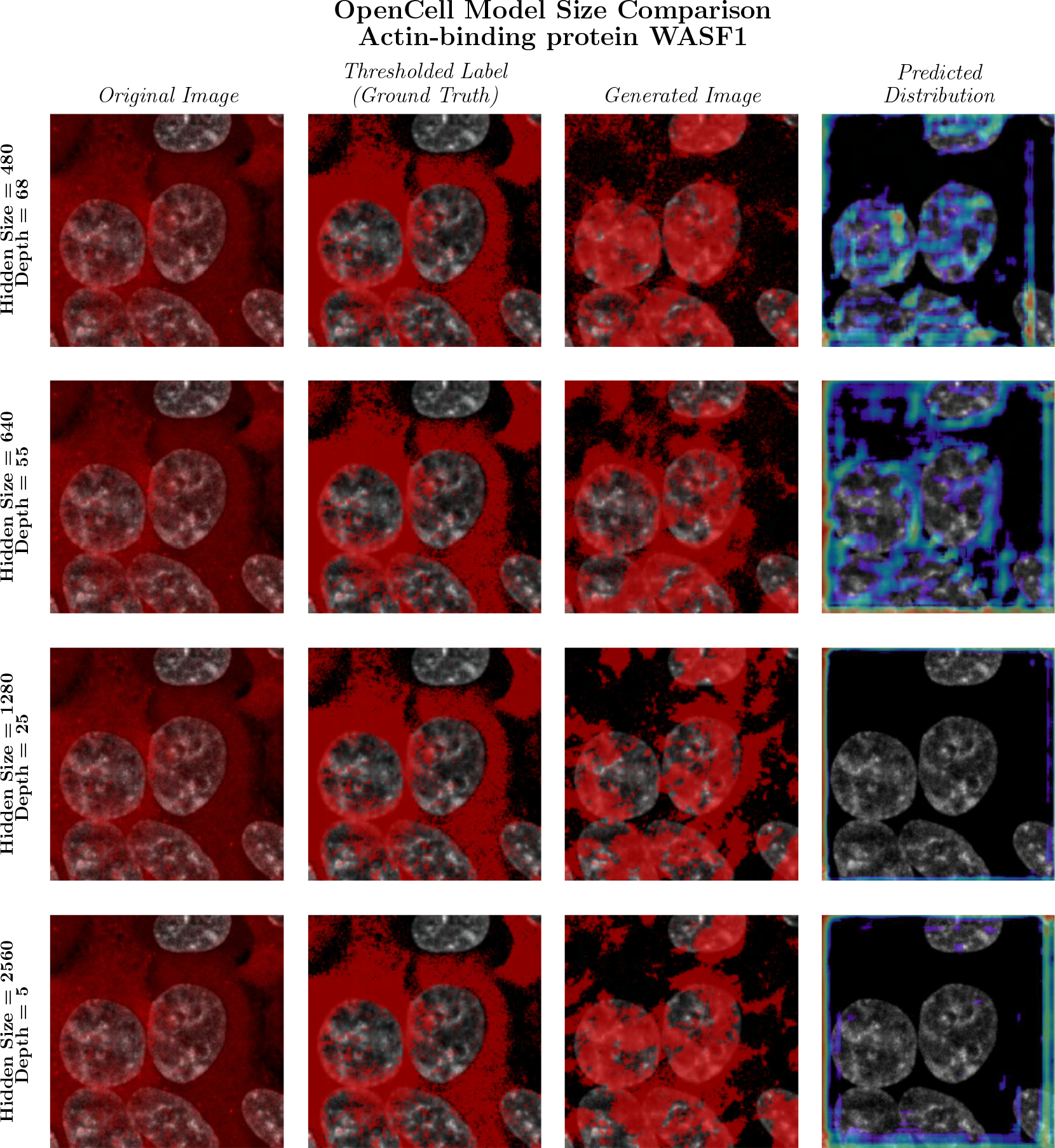
Similar to Fig. S6, we depict the performance of CELL-E 2 models only trained on the OpenCell dataset. We see the best performance on the 480 model, but not drastically different predicted distribution images. This is likely a function of reduced training time due to the quick overfitting of the model.

### C.3 Finetuning

Table S8 shows the image prediction performance of models across datasets after fine-tuning on the OpenCell dataset. Table S9 shows the sequence prediction accuracy of the same models.

**Table S5:** ESM-2 Masked Sequence In-Filling Accuracy (No Image)

| Training Set Proteins |  |  |  |  |
| --- | --- | --- | --- | --- |
| Dataset | Hidden Size | # Layers | Sequence MAE | Cosine Similarity |
| HPA | 480 | 12 | .7351 ± .1100 | .9464 ± .0232 |
| | 640 | 30 | .6507 ± .1317 | .9572 ± .0183 |
| | 1280 | 33 | .4921 ± .1741 | .9724 ± .0133 |
|  | 2560 | 36 | **.3818 ± .1911** | **.9778 ± .0130** |
| OpenCell | 480 | 12 | .7276 ± .1144 | .9425 ± .0233 |
| | 640 | 30 | .6151 ± .1364 | .9572 ± .0159 |
| | 1280 | 33 | .4335 ± .1650 | .9746 ± .0082 |
|  | 2560 | 36 | **.3298 ± .1762** | **.9793 ± .0089** |
| Validation Set Proteins |  |  |  |  |
| Dataset | Hidden Size | # Layers | Sequence MAE | Cosine Similarity |
| HPA | 480 | 12 | .7368 ± .1116 | .9471 ± .0209 |
| | 640 | 30 | .6553 ± .1334 | .9571 ± .0161 |
| | 1280 | 33 | .5005 ± .1705 | .9723 ± .0096 |
|  | 2560 | 36 | **.3894 ± .1911** | **.9777 ± .0096** |
| OpenCell | 480 | 12 | .7355 ± .1130 | .9381 ± .0286 |
| | 640 | 30 | .6185 ± .1454 | .9538 ± .0199 |
| | 1280 | 33 | .4260 ± .1822 | .9737 ± .0096 |
|  | 2560 | 36 | **.3220 ± .1848** | **.9789 ± .0086** |

## D Discussion

### D.1 CELL-E Comparison

Table S10 shows the image prediction metrics for the original CELL-E model on both the HPA and OpenCell datasets. Note that CELL-E was only trained on OpenCell data.

Table S11 depicts the mean time taken for 10 separate model predictions. CELL-E is not directly comparable to CELL-E 2 due to differences in language model and package versioning, so we opt to include the compute time of CELL-E 2 using an autoregressive reconstruction scheme (i.e. 256 sequential steps from top left to bottom right). CELL-E 2 model run in autoregressive mode are significantly slower due to the lack of cache implementation found in CELL-E and the larger ESM-2 language model compared to the TAPE model used in CELL-E. CELL-E 2 models which generate the prediction in a single step (NAR) are an orders of magnitude faster than their autoregressive counterparts.

### D.2 De novo NLS Design

**NLS generation**

1. We selected a desired NLS length (iterating over a range of 5 to 30 residues) and inserted that number of mask tokens after the starting methionine in the GFP sequence. (e.g. an NLS of length 5 at the N terminus would have an input sequence of <START> M <MASK_SEQ> <MASK_SEQ> <MASK_SEQ> <MASK_SEQ> <MASK_SEQ>SKGEE…<END> <PAD>…).
2. We randomly chose a nucleus image and segmented the nuclei area by applying a mask with Cellpose [14]. We assigned the pixels inside the nucleus area to True and used this as the threshold image.
3. We inputted the masked GFP sequence, the nucleus image, and the threshold image to the transformer and sampled the output. We used the model depth that achieved the highest performance on sequence reconstruction, which was `OpenCell_2560`.

**Table S6:** Masked Sequence In-Filling Accuracy.
4. For each sequence length, we generated 300 candidates per length per terminus. We then provided the `HPA Finetuned (Finetuned HPA VQGAN)_480` model with the predicted NLS + GFP sequence and the nucleus image. Using the previously calculated nucleus mask, we calculate the percentage of positive intensity predicted within the nucleus bounds. Any sequence with a predicted nucleus proportion intensity *<* 75% was discarded.

| Training Set Proteins |  |  |  |  |  |
| --- | --- | --- | --- | --- | --- |
| Dataset | Train Set | Hidden Size | Depth | Sequence MAE | Cosine Similarity |
| HPA | HPA | 480 | 68 | .8548 ± .1050 | .9500 ± .0260 |
| | | 640 | 55 | .7738 ± .1368 | .9580 ± .0238 |
| | | 1280 | 25 | .5818 ± .2053 | **.9733 ± .0195** |
| | | 2560 | 5 | **.5294 ± .2402** | .9732 ± .0235 |
| | OpenCell | 480 | 68 | .8554 ± .1047 | .9504 ± .0262 |
| | | 640 | 55 | .7806 ± .1343 | .9576 ± .0239 |
| | | 1280 | 25 | .6377 ± .1850 | .9709 ± .0191 |
| | | 2560 | 5 | .5599 ± .2294 | .9721 ± .0235 |
| OpenCell | HPA | 480 | 68 | .8403 ± .1102 | .9463 ± .0277 |
| | | 640 | 55 | .7434 ± .1356 | .9557 ± .0263 |
| | | 1280 | 25 | .5315 ± .1996 | .9725 ± .0219 |
| | | 2560 | 5 | .4760 ± .2281 | **.9726 ± .0266** |
| | OpenCell | 480 | 68 | .7507 ± .1709 | .9533 ± .0285 |
| | | 640 | 55 | .6641 ± .1764 | .9610 ± .0272 |
| | | 1280 | 25 | .5698 ± .2016 | .9709 ± .0220 |
| | | 2560 | 5 | **.4950 ± .2456** | .9711 ± .0271 |
| Validation Set Proteins |  |  |  |  |  |
| Dataset | Train Set | Hidden Size | Depth | Sequence MAE | Cosine Similarity |
| HPA | HPA | 480 | 68 | .8628 ± .0951 | .9504 ± .0237 |
| | | 640 | 55 | .7917 ± .1245 | .9577 ± .0216 |
| | | 1280 | 25 | .6512 ± .1794 | .9708 ± .0163 |
| | | 2560 | 5 | .5759 ± .2322 | .9722 ± .0210 |
| | OpenCell | 480 | 68 | .8625 ± .0935 | .9508 ± .0240 |
| | | 640 | 55 | .7927 ± .1245 | .9577 ± .0216 |
| | | 1280 | 25 | .6476 ± .1811 | .9711 ± .0163 |
|  |  | 2560 | 5 | **.5696 ± .2288** | **.9724 ± .0210** |
| OpenCell | HPA | 480 | 68 | .8651 ± .0992 | .9420 ± .0312 |
| | | 640 | 55 | .7675 ± .1318 | .9529 ± .0271 |
| | | 1280 | 25 | .5910 ± .2065 | .9699 ± .0213 |
| | | 2560 | 5 | .5137 ± .2414 | .9700 ± .0250 |
| | OpenCell | 480 | 68 | .8600 ± .1030 | .9430 ± .0316 |
| | | 640 | 55 | .7645 ± .1332 | .9532 ± .0273 |
| | | 1280 | 25 | .5872 ± .2060 | .9703 ± .0213 |
|  |  | 2560 | 5 | **.5080 ± .2365** | **.9703 ± .0250** |

We generated candidate NLS with lengths from 2 to 30 amino acids at the N and C termini of the protein. We ranked them using these criteria:

- Forward Consistency: The proportion of positive signal in the nucleus mask relative to the whole image, using the best image prediction model (480 model), similar to Section 5.1.

**Table S7:** Masked Sequence Random In-Filling Accuracy.

| Training Set Proteins |  |  |
| --- | --- | --- |
| Dataset | Sequence MAE | Cosine Similarity |
| HPA | .9600 ± .0274 | .9502 ± .0181 |
| OpenCell | .9603 ± .0268 | .9469 ± .0176 |
| Validation Set Proteins |  |  |
| Dataset | Sequence MAE | Cosine Similarity |
| HPA | .9605 ± .0257 | .9509 ± .0157 |
| OpenCell | .9592 ± .0282 | .9461 ± .0191 |

**Figure S8:**
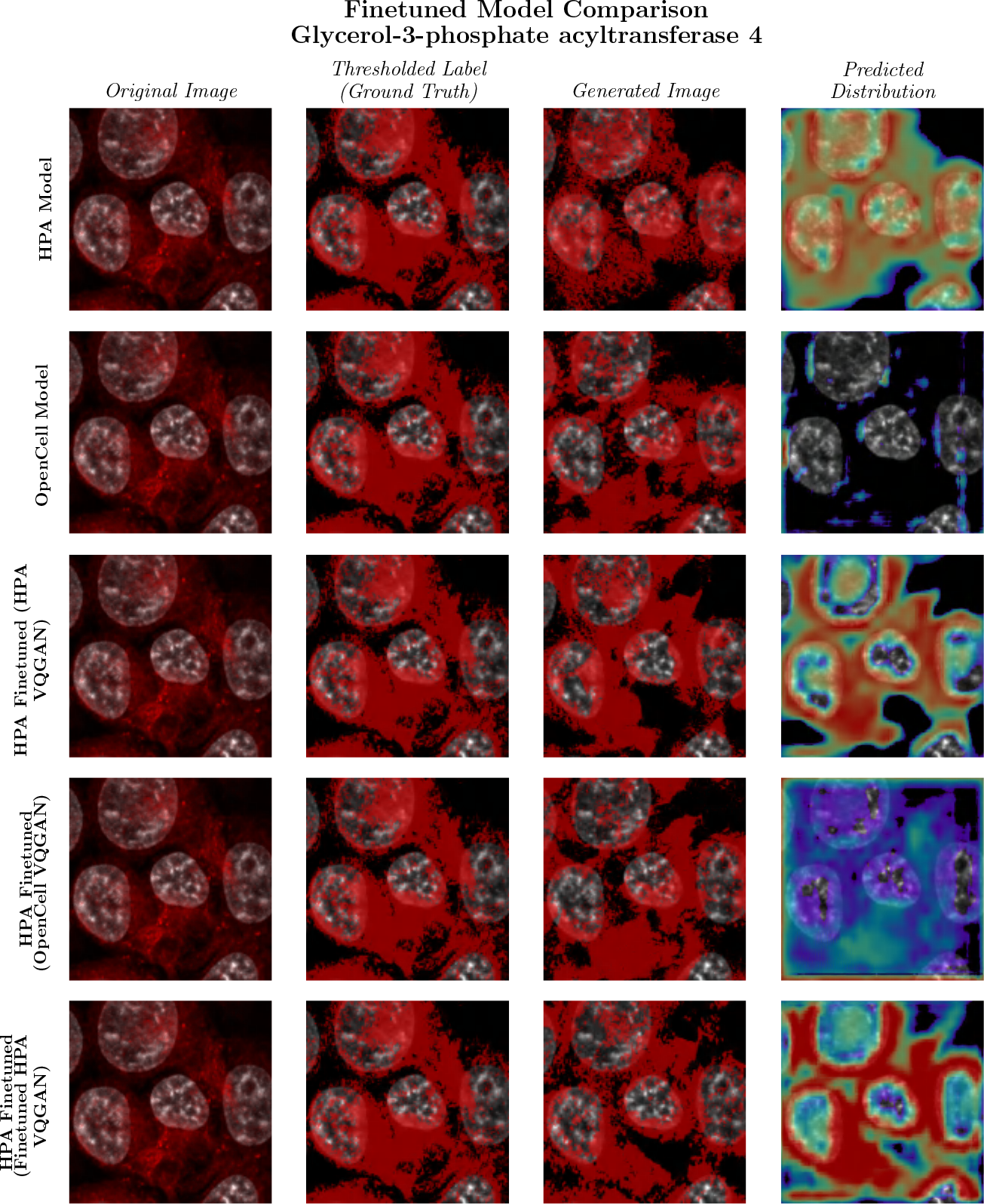
Various model performance from different fine tuning methods. We note superior predictive performance from the model with where we initially fine-tune the image encoder.
- Image Prediction Confidence: The values from the predicted distribution using a masked approach, indicating the confidence in the localization image prediction.

**Table S8:** Image Prediction Accuracy after Finetuning on HPA and OpenCell.
- Text Prediction Confidence: The average probability values of the predicted NLS sequence tokens.
- Sequence Similarity: The maximum alignment score between the candidate NLS and sequences from the NLSdb, similar to Madani et. al. [15].
- Embedding Cosine Angle: The minimum cosine angle between the embeddings of the candidate NLS and sequences from the NLdb [16], using the same language model from Section 5.2, except similarity is evaluated on the entire protein sequence (NLS + GFP), rather than limited to the masked positions.

| Training Set Proteins |  |  |  |  |  |  |  |  |  |
| --- | --- | --- | --- | --- | --- | --- | --- | --- | --- |
| Dataset | Image Encoders | Hidden Size | Depth | Nucleus Proportion MAPE | Image MAE | PDF MAE | SSIM | FID | IS |
| HPA | HPA | 480 | 68 | .0292 ± .0291 | .3606 ± .0832 | .3599 ± .0836 | **.2237 ± .1479** | 22.0947 | 2.8130 ± .0208 |
|  | OpenCell |  |  | **.0245 ± .0317** | .4680 ± .0776 | .3428 ± .0833 | .1047 ± .0840 | 23.2398 | 3.0922 ± .0167 |
|  | HPA Finetuned |  |  | .0249 ± .0289 | .3755 ± .1011 | **.3292 ± .0848** | .1406 ± .1027 | 8.3675 | **3.9647 ± .0299** |
|  | HPA | 640 | 55 | .0299 ± .0263 | .3475 ± .0834 | .3472 ± .0819 | .1516 ± .1118 | 6.7563 | 2.0455 ± .0099 |
|  | OpenCell |  |  | .0273 ± .0254 | .4518 ± .0570 | .3505 ± .0778 | .0900 ± .0747 | 31.7937 | 2.5763 ± .0119 |
|  | HPA Finetuned |  |  | .0270 ± .0249 | **.3041 ± .0907** | .3328 ± .0794 | .1278 ± .0910 | 11.4788 | 2.3392 ± .0130 |
|  | HPA | 1280 | 25 | .0448 ± .0400 | .3461 ± .0820 | .3350 ± .0842 | .2004 ± .1364 | 6.8770 | 2.1677 ± .0096 |
|  | OpenCell |  |  | .0426 ± .0410 | .4486 ± .0556 | .3401 ± .0826 | .1067 ± .0841 | 17.6565 | 2.7158 ± .0105 |
|  | HPA Finetuned |  |  | .0435 ± .0437 | .3315 ± .0888 | .3323 ± .0826 | .1762 ± .1183 | **5.9633** | 2.2360 ± .0279 |
|  | HPA | 2560 | 5 | .0729 ± .0655 | .3844 ± .0704 | .3590 ± .0792 | .1793 ± .1161 | 12.6113 | 2.0646 ± .0112 |
|  | OpenCell |  |  | .0727 ± .0776 | .4736 ± .0633 | .3428 ± .0847 | .1291 ± .0925 | 8.4963 | 2.1803 ± .0116 |
|  | HPA Finetuned |  |  | .0744 ± .0671 | .3507 ± .0803 | .3599 ± .0795 | .2014 ± .1322 | 16.672 | 2.2908 ± .0156 |
| OpenCell | HPA | 480 | 68 | .0157 ± .0151 | .3712 ± .0791 | .3699 ± .0799 | .2038 ± .1525 | 17.1616 | 3.0822 ± .0843 |
|  | OpenCell |  |  | **.0135 ± .0135** | .4996 ± .0007 | **.3161 ± .1117** | .1874 ± .1495 | 1.5167 | **3.0898 ± .1459** |
|  | HPA Finetuned |  |  | .0154 ± .0150 | **.3170 ± .1159** | .3186 ± .1215 | **.2125 ± .1600** | 18.7426 | 3.9276 ± .1406 |
|  | HPA | 640 | 55 | .0165 ± .0151 | .4011 ± .0667 | .3439 ± .1026 | .1263 ± .1063 | 6.0163 | 2.2918 ± .0533 |
|  | OpenCell |  |  | .0149 ± .0136 | .4732 ± .0192 | .3415 ± .1054 | .1356 ± .1281 | 4.9600 | 2.4016 ± .0866 |
|  | HPA Finetuned |  |  | .0167 ± .0150 | .3305 ± .1035 | .3400 ± .1059 | .1525 ± .1195 | 2.8065 | 2.7464 ± .0621 |
|  | HPA | 1280 | 25 | .0243 ± .0224 | .3817 ± .0686 | .3355 ± .1065 | .1546 ± .1201 | 3.7530 | 2.5043 ± .0454 |
|  | OpenCell |  |  | .0220 ± .0205 | .4671 ± .0278 | .3236 ± .1089 | .1702 ± .1491 | **.5084** | 3.0222 ± .1054 |
|  | HPA Finetuned |  |  | .0254 ± .0241 | .3701 ± .0838 | .3581 ± .1054 | .1468 ± .1156 | 5.2415 | 2.5990 ± .1403 |
|  | HPA | 2560 | 5 | .0411 ± .0379 | .4067 ± .0745 | .3363 ± .1087 | .1775 ± .1299 | 14.7029 | 2.4132 ± .0603 |
|  | OpenCell |  |  | .0540 ± .0492 | .4977 ± .0124 | .3753 ± .1089 | .1630 ± .1200 | 26.8886 | 1.8080 ± .0489 |
|  | HPA Finetuned |  |  | .0394 ± .0359 | .3710 ± .0843 | .3492 ± .1032 | .1727 ± .1265 | 15.3433 | 2.5426 ± .0637 |
| Validation Set Proteins |  |  |  |  |  |  |  |  |  |
| Dataset | Image Encoders | Hidden Size | Depth | Nucleus Proportion MAPE | Image MAE | PDF MAE | SSIM | FID | IS |
| HPA | HPA | 480 | 68 | .0291 ± .0259 | .3589 ± .0838 | .3583 ± .0843 | **.2246 ± .1501** | 21.8254 | 2.8176 ± .0210 |
|  | OpenCell |  |  | **.0245 ± .0233** | .4681 ± .0774 | .3430 ± .0833 | .1047 ± .0853 | 23.9367 | 3.0918 ± .0519 |
|  | HPA Finetuned |  |  | .0249 ± .0235 | .3427 ± .0908 | **.3292 ± .0847** | .1397 ± .1047 | 8.7002 | **3.9302 ± .0716** |
|  | HPA | 640 | 55 | .0304 ± .0273 | .3469 ± .0835 | .3476 ± .0821 | .1496 ± .1117 | 7.0875 | 2.0259 ± .0310 |
|  | OpenCell |  |  | .0276 ± .0265 | .4519 ± .0567 | .3502 ± .0779 | .0905 ± .0759 | 31.8870 | 2.5738 ± .0402 |
|  | HPA Finetuned |  |  | .0279 ± .0262 | **.3041 ± .0906** | .3326 ± .0793 | .1266 ± .0917 | 12.0062 | 2.3105 ± .0310 |
|  | HPA | 1280 | 25 | .0454 ± .0434 | .3462 ± .0822 | .3362 ± .0847 | .1984 ± .1368 | 6.8893 | 2.1656 ± .0288 |
|  | OpenCell |  |  | .0433 ± .0444 | .4484 ± .0560 | .3400 ± .0827 | .1064 ± .0848 | 18.1654 | 2.7017 ± .0460 |
|  | HPA Finetuned |  |  | .0430 ± .0403 | .3322 ± .0882 | .3320 ± .0824 | .1771 ± .1162 | **5.9752** | 2.2687 ± .0112 |
|  | HPA | 2560 | 5 | .0746 ± .0686 | .3828 ± .0708 | .3594 ± .0807 | .1790 ± .1176 | 12.6199 | 2.0311 ± .0311 |
|  | OpenCell |  |  | .0739 ± .0755 | .4730 ± .0650 | .3429 ± .0854 | .1289 ± .0957 | 8.7266 | 2.1980 ± .0275 |
|  | HPA Finetuned |  |  | .0761 ± .0697 | .3510 ± .0816 | .3603 ± .0810 | .2003 ± .1332 | 16.4098 | 2.2785 ± .0319 |
| OpenCell | HPA | 480 | 68 | .0166 ± .0151 | .3776 ± .0834 | .3477 ± .1268 | **.1869 ± .1503** | 17.4075 | 2.9113 ± .1199 |
|  | OpenCell |  |  | **.0159 ± .0156** | .4996 ± .0006 | .3506 ± .1208 | .1574 ± .1372 | 2.5026 | 2.7168 ± .1137 |
|  | HPA Finetuned |  |  | .0170 ± .0160 | **.3449 ± .1305** | .3487 ± .1340 | .1881 ± .1541 | 19.2683 | **3.6083 ± .2013** |
|  | HPA | 640 | 55 | .0176 ± .0155 | .4028 ± .0668 | .3644 ± .1004 | .1060 ± .0928 | 7.9330 | 2.0560 ± .1219 |
|  | OpenCell |  |  | .0170 ± .0149 | .4771 ± .0201 | .3684 ± .1073 | .1081 ± .1121 | 5.1479 | 2.1141 ± .1304 |
|  | HPA Finetuned |  |  | .0172 ± .0151 | .3477 ± .1043 | .3583 ± .1033 | .1339 ± .1083 | 2.4811 | 2.4813 ± .1009 |
|  | HPA | 1280 | 25 | .0258 ± .0243 | .3890 ± .0709 | .3572 ± .1050 | .1355 ± .1092 | 3.7844 | 2.2680 ± .1109 |
|  | OpenCell |  |  | .0262 ± .0259 | .4743 ± .0275 | .3576 ± .1133 | .1339 ± .1218 | **.9963** | 2.6376 ± .1468 |
|  | HPA Finetuned |  |  | .0247 ± .0234 | .3599 ± .0813 | **.3361 ± .1078** | .1645 ± .1229 | 4.8118 | 2.8837 ± .0426 |
|  | HPA | 2560 | 5 | .0464 ± .0464 | .4081 ± .0776 | .3591 ± .1074 | .1598 ± .1211 | 13.7206 | 2.2251 ± .1164 |
|  | OpenCell |  |  | .0594 ± .0533 | .4969 ± .0121 | .3928 ± .1074 | .1509 ± .1135 | 27.7841 | 1.7532 ± .0837 |
|  | HPA Finetuned |  |  | .0446 ± .0430 | .3812 ± .0885 | .3709 ± .0988 | .1549 ± .1193 | 13.4599 | 2.3191 ± .1147 |

We rounded all values to one decimal place and ranked them by 1) Sequence Similarity, 2) Embedding Cosine Similarity, 3) Forward Consistency, 4) Image Prediction Confidence, 5) Text Prediction Confidence.

**Table S9:**
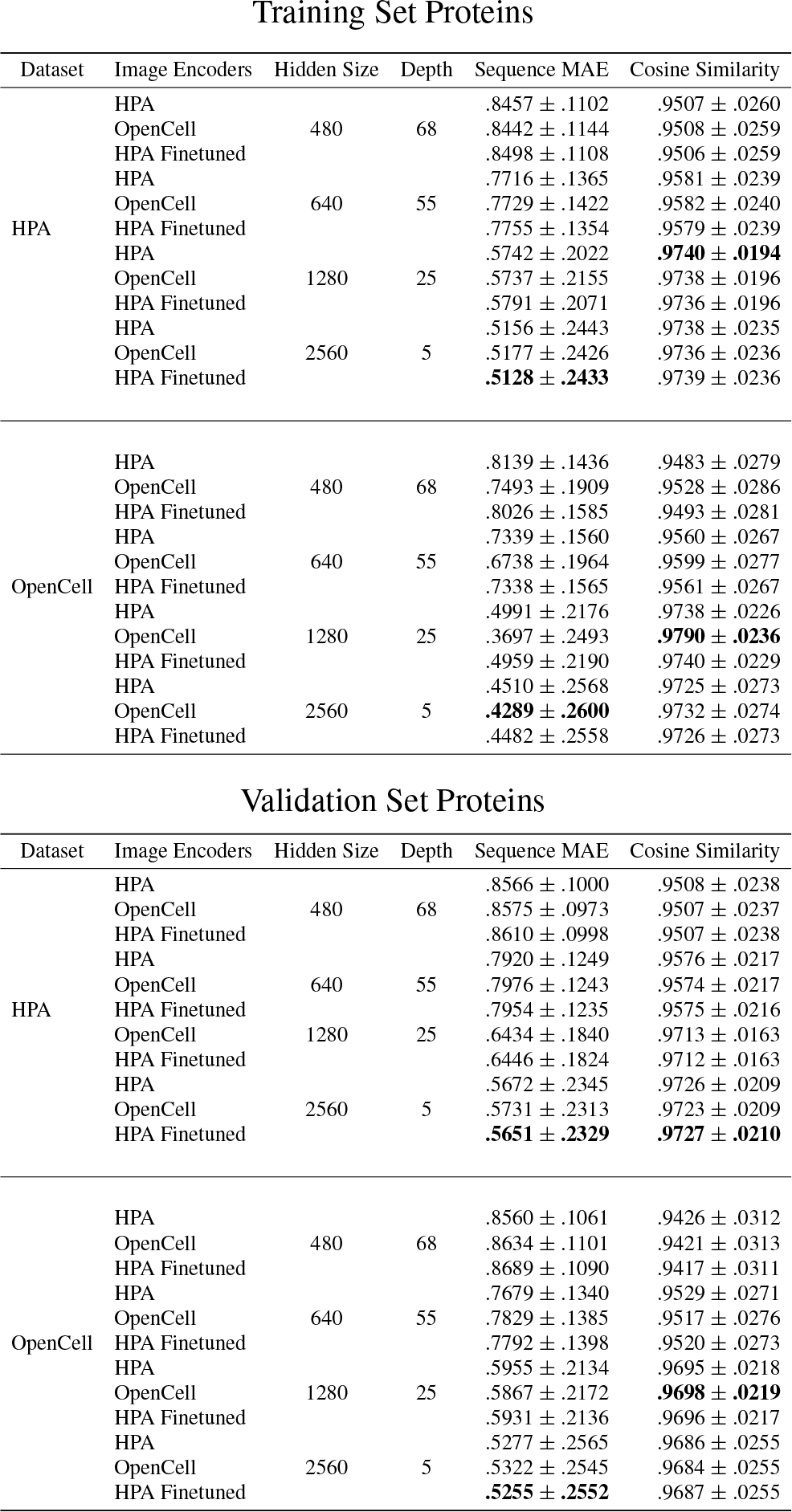
Masked Sequence In-Filling Accuracy after Finetuning on HPA and OpenCell.

**Table S10:** Image Prediction Accuracy for CELL-E.

| Training Set Proteins |  |  |  |  |  |  |  |  |
| --- | --- | --- | --- | --- | --- | --- | --- | --- |
| Dataset | Hidden Size | Depth | Nucleus Proportion MAPE | Image MAE | PDF MAE | SSIM | FID | IS |
| HPA | 768 | 32 | .0672 ± .0632 | .3601 ± .0829 | .3303 ± .0796 | .2219 ± .1383 | 4.2355 | 2.1292 ± .0139 |
| OpenCell | | | .0377 ± .0327 | .3642 ± .1150 | .3600 ± .1044 | .2133 ± .1825 | 11.3911 | 2.4390 ± .0625 |
| Validation Set Proteins |  |  |  |  |  |  |  |  |
| Dataset | Hidden Size | Depth | Nucleus Proportion MAPE | Image MAE | PDF MAE | SSIM | FID | IS |
| HPA | 768 | 32 | .0786 ± .0644 | .3610 ± .0816 | .3308 ± .0785 | .2217 ± .1371 | 4.1685 | 2.1021 ± .0371 |
| OpenCell | | | .0347 ± .0294 | .3671 ± .1117 | .3653 ± .1008 | .2060 ± .1846 | 10.5555 | 2.4762 ± .0866 |

**Table S11:** Speed Comparison.

| Model | Hidden Size | Autoregressive | Mean Generation Time (s) |
| --- | --- | --- | --- |
| CELL-E (Cached) | 768 | Yes | $18.2740 \pm 0.0451$ |
| CELL-E (Non-Cached) | 768 | Yes | $28.7694 \pm 0.3207$ |
| CELL-E 2 | 480 | Yes | $55.0057 \pm 0.2069$ |
| CELL-E 2 | 640 | Yes | $62.9650 \pm 0.1033$ |
| CELL-E 2 | 1280 | Yes | $74.3698 \pm 0.1788$ |
| CELL-E 2 | 2560 | Yes | $128.9960 \pm 0.3718$ |
| **CELL-E 2** | **480** | **No** | **$0.2784 \pm 0.0006$** |
| CELL-E 2 | 640 | No | $0.3067 \pm 0.0012$ |
| CELL-E 2 | 1280 | No | $0.3249 \pm 0.0011$ |
| CELL-E 2 | 2560 | No | $0.5487 \pm 0.0022$ |

**Table S12:** NLS Composition.

| Max ID % | # Sequences | Mean Sequence Length | Mean % R or K |
| --- | --- | --- | --- |
| 0% - 33% | 109 | $25.6606 \pm 3.0099$ | $20.6379 \pm 8.6101$ |
| 33% - 66% | 133 | $17.1955 \pm 5.0804$ | $32.0076 \pm 12.8334$ |
| 66% - 100% | 13 | $6.9231 \pm 1.2558$ | $57.5794 \pm 17.9351$ |

### D.3 Visualizing Attention

In Fig. S11 and Fig. 3, we depict the relative attention weights placed on the input amino acid sequence and nucleus image used to generate the threshold prediction. Specifically, we sought to emphasize weights correlated with positive signal, that is patches with largely white pixels. In this way, we do not bias the weights we consider with the use of any manual feature annotations or image segmentation. We first use attention rollout [17] to obtain the relative correlation between tokens at the end of the network. We then take an average across the multiplied attention heads. From here, we separate “positive” vs “negative” signal image patches based on the average intensity within the predicted image. Positive and negative patches are those where ≥ 75% and ≤ 25% are white, respectively. We then subtract the mean attention weights of the negative patches from the positive patches. Those with positive differences are therefore more correlated with a positive signal prediction in the cell. For visualization, we depict the log value of the difference (normalized to 1).

Values used to sort candidate NLS sequences are available in the `de_novo_NLS_sequences.csv`.

Predicted sequences are shown in Table S13.

**Figure S9:**
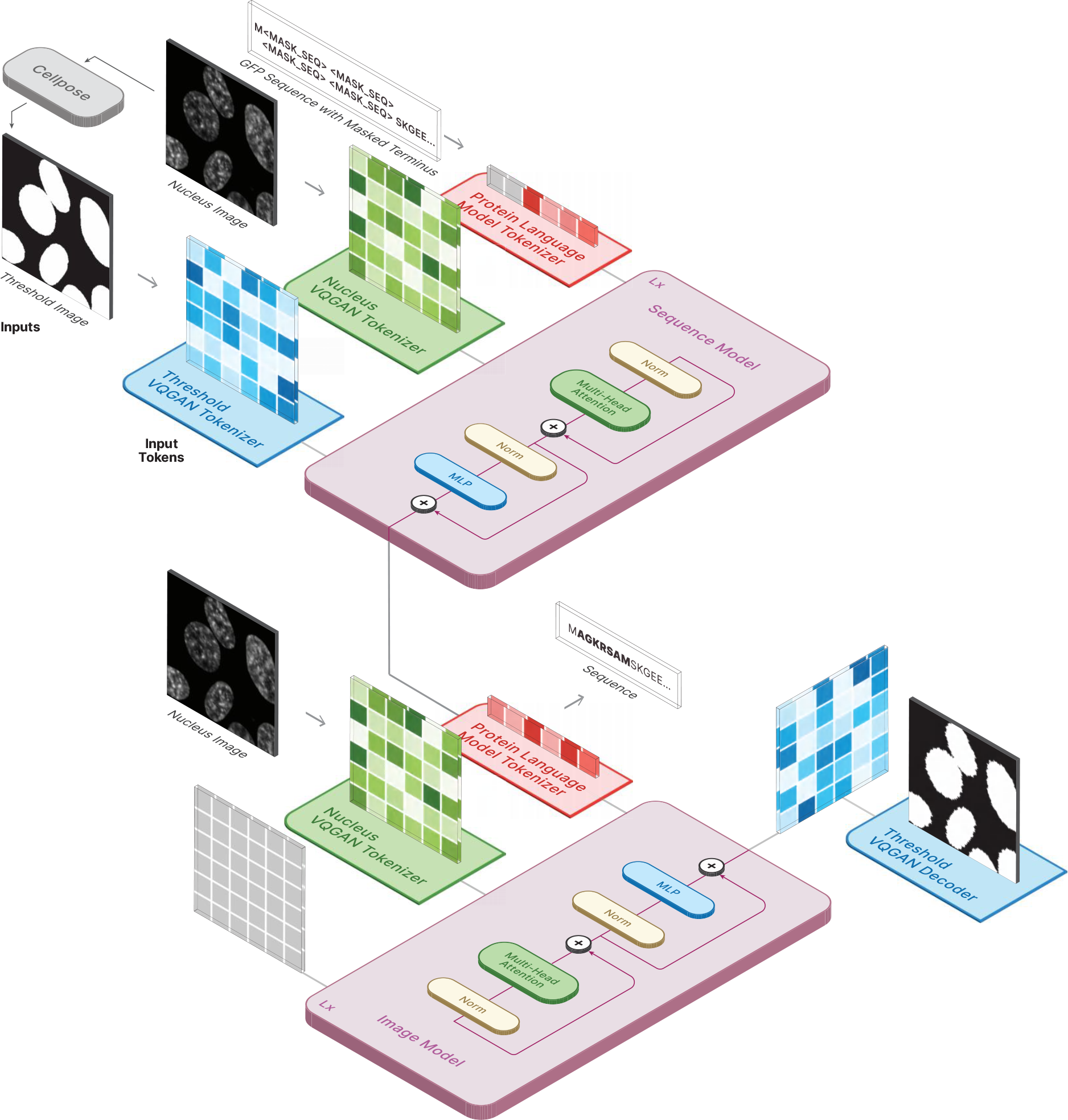
Diagram depicts the pipeline for NLS discovery. In the top half, we predetermine the length of the novel NLS sequence and insert the corresponding number of mask tokens either after the starting Methionine or before the `<END>` token, depending on the chosen terminus. The threshold image is obtained by passing the nucleus image through Cellpose. In the bottom half, we pass the the GFP with proposed NLS sequence into an image prediction model to ensure predictive consistency of the sequence.

**Figure S10:**
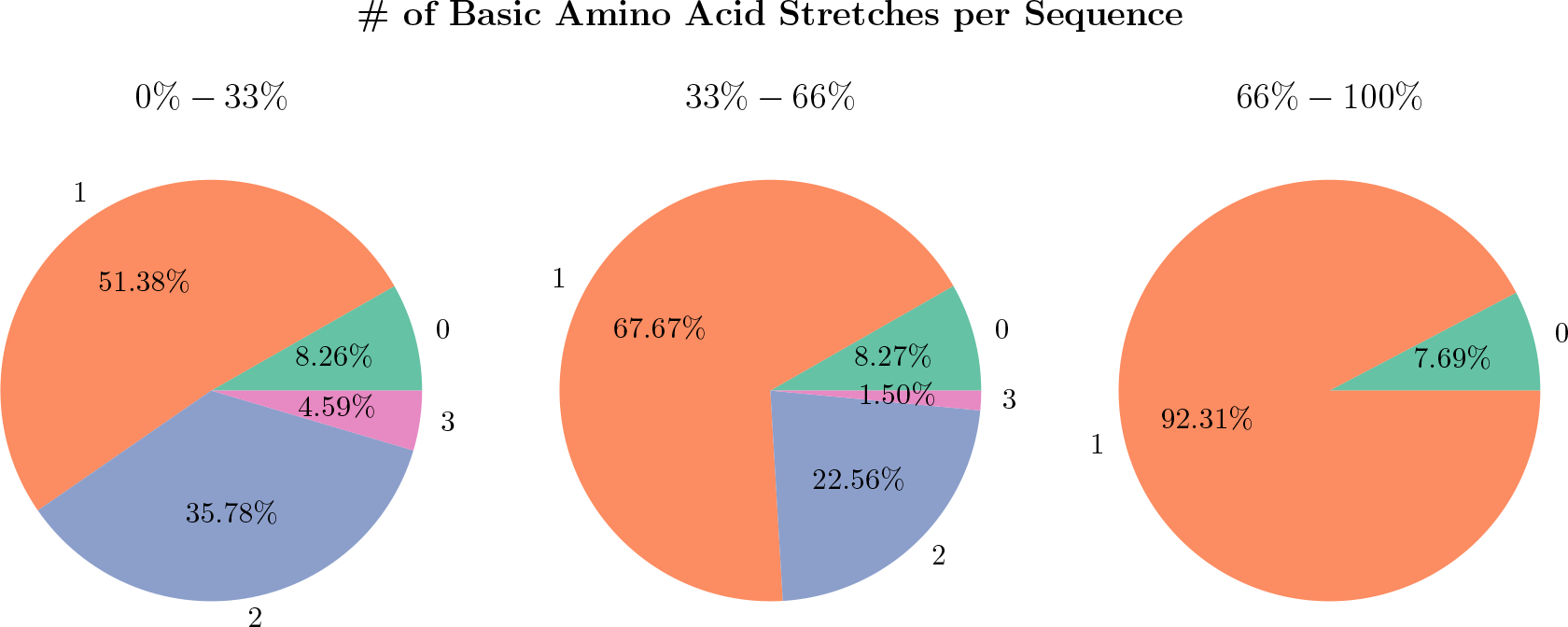
Pie charts showing the maximum # of stretches (numbers outside of circle) of R and K amino acids per proposed NLS sequence. Stretches are calculated based on the number of continuous R and K amino acids with a maximum tolerance of 2 amino acid gap. Only streches with 4 or more amino acids are counted. Proteins are shown binned with respect to Max ID % sequence homology with the NLSdb (0%-33%, 33%-66%, and 66%-100%). The relative proportion of max stretches per bin is shown as a percentage inside the circle.

**Figure S11:**
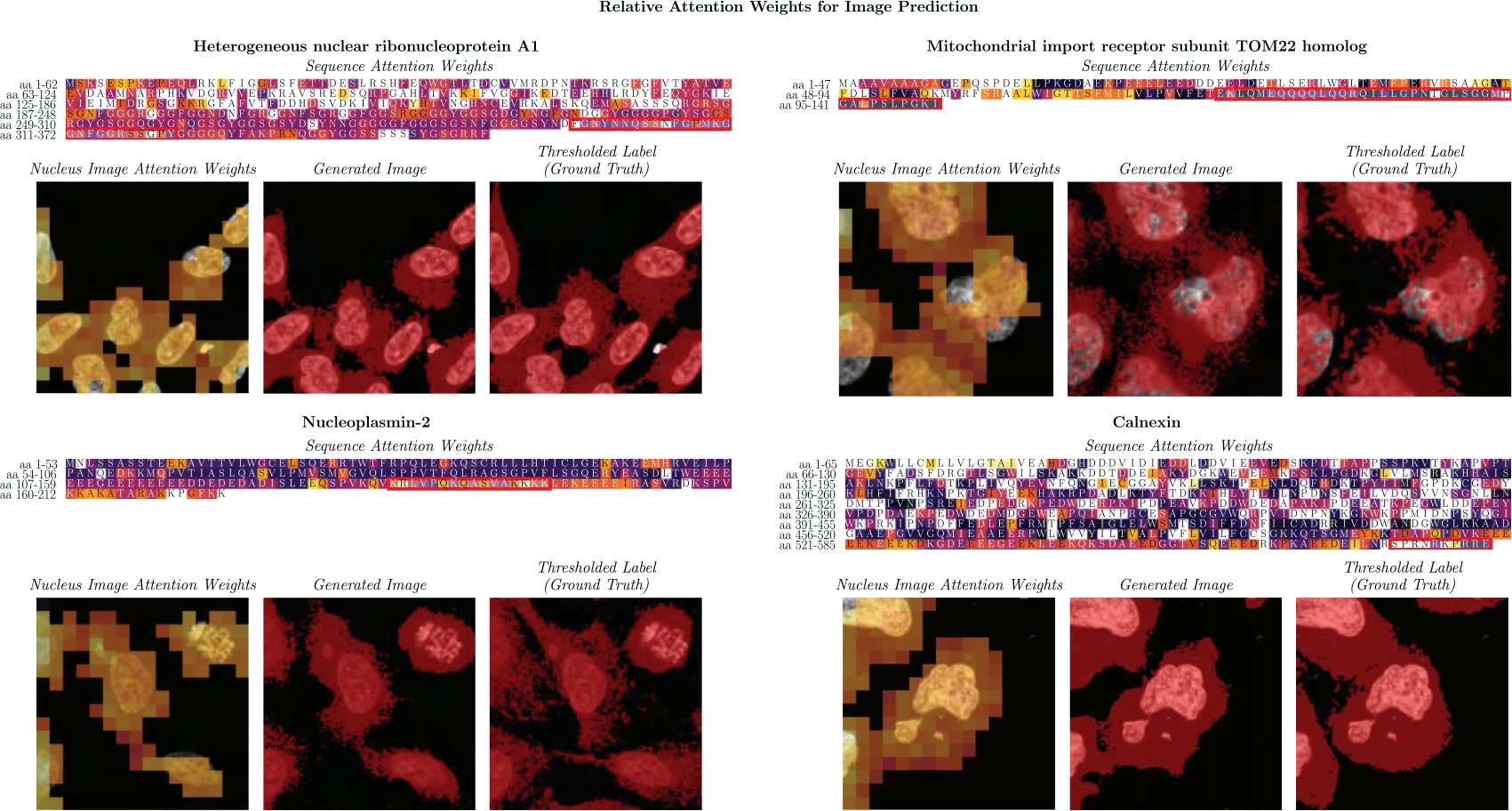
Relative attention weights of predictions from `HPA_480` on HPA images with known localization signals (highlighted in red). Three proteins with documented localization signals show high attention on those regions: Heterogeneous nuclear riboprotein A1 (top left), which localizes to the nucleus and cytoplasm [18, 19]; Nucleoplasmin-2 (bottom left), which localizes to the nucleus [20]; and Mitochondrial import receptor subunit TOM22 homolog (top right), which localizes to the mitochondria [21]. However, Calnexin (bottom right), which localizes to the endoplasmic reticulum [22], does not show high attention on its localization signal despite the correct prediction. This may be due to the loss of subcellular features in the thresholding process caused by the low resolution of the fluorescence image. We also observe high attention on other amino acids in the sequences that are not known localization signals. These may indicate potential sites of interest for further biological investigation.

**Table S13:**
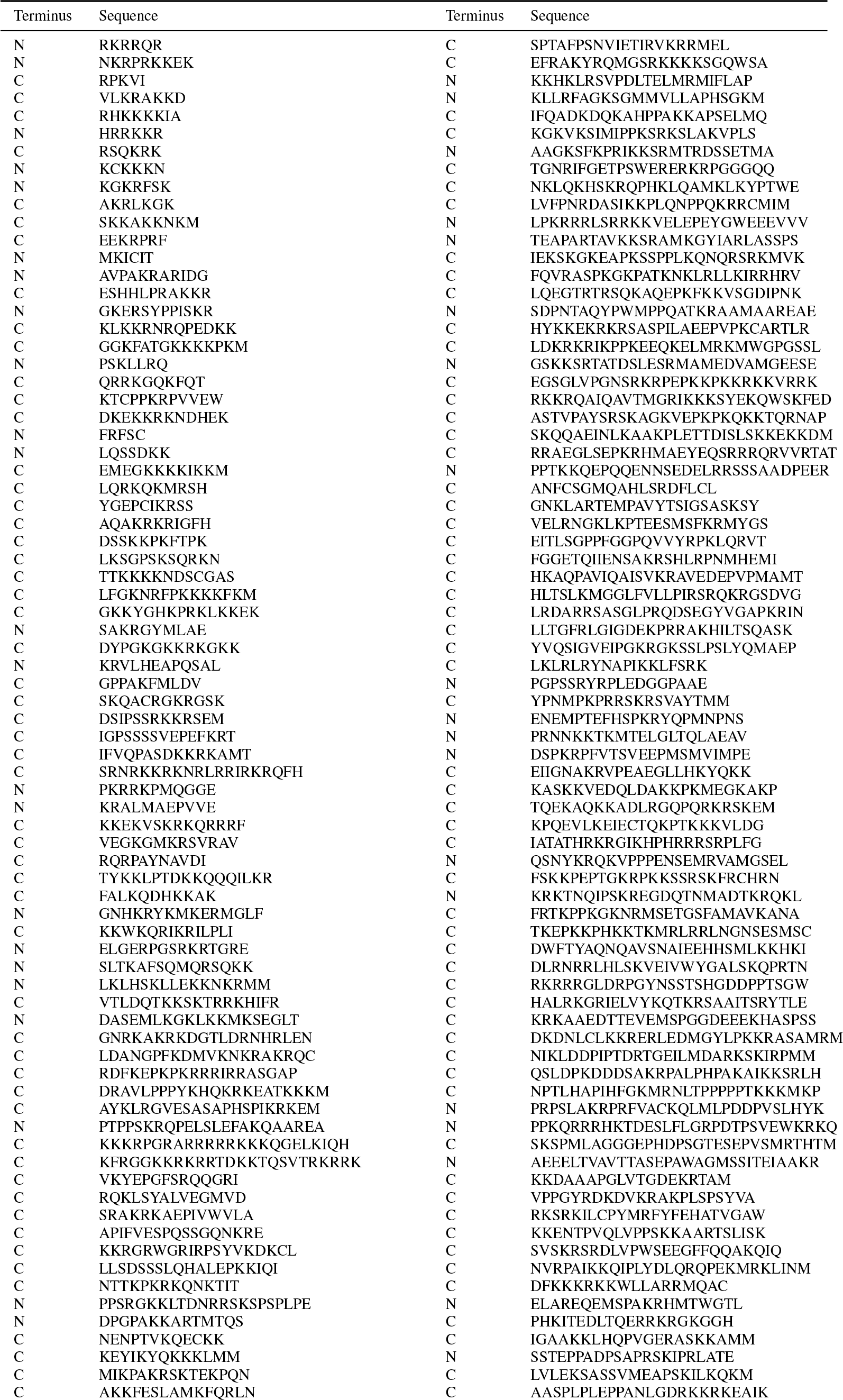
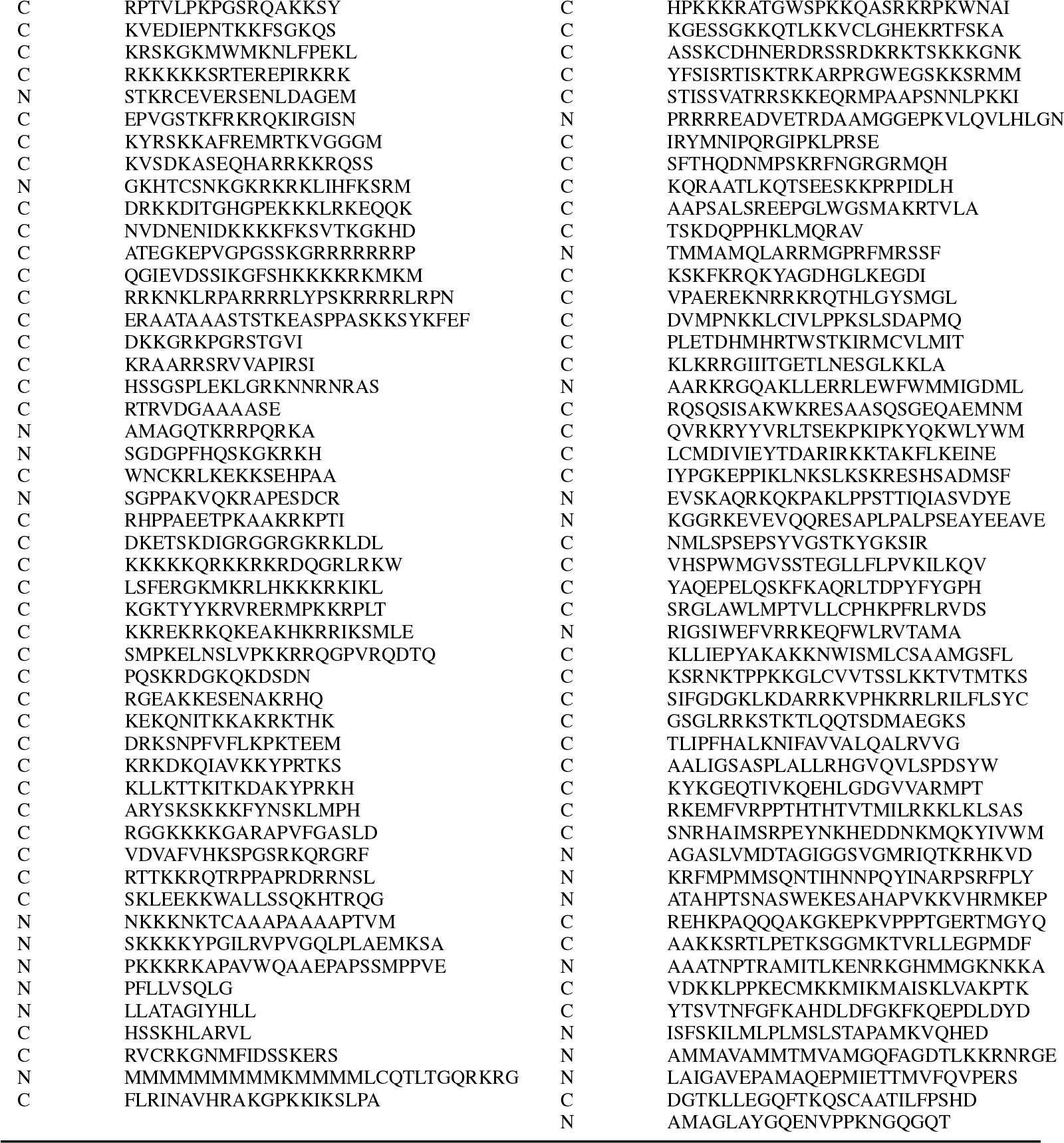
NLS candidates sorted by nucleus proportion.

